# RNA viruses that exploit self-cleaving ribozymes for translation initiation

**DOI:** 10.1101/2024.05.16.594327

**Authors:** María José López-Galiano, Olga Rueda, Sotaro Chiba, Marco Forgia, Beatriz Navarro, Amelia Cervera, Artem Babaian, Francesco Di Serio, Massimo Turina, Marcos de la Peña

## Abstract

Small self-cleaving ribozymes are catalytic RNAs originally discovered in viroid-like agents, which are replicating circular RNAs (circRNAs) postulated as relics of a prebiotic RNA world. In the last decade, however, small ribozymes have also been detected across the tree of life, from bacterial to human genomes, and more recently, in unusual circRNA viruses. Here we report the conserved occurrence of diverse small ribozymes within the linear genomes of typical double- and single-stranded RNA virus families from fungi and plants. Type I hammerhead ribozyme motifs occur in the 5’-UTR regions of chrysovirids and fusarivirids, displaying self-cleaving activity *in vitro* and *in vivo*. Similar hammerhead, as well as hepatitis delta and twister ribozymes, are also found in diverse megabirna-, hypo-, fusagra-, toti-or tombus-like viruses among others. The ribozymes occur not only as isolated motifs within UTRs but also as tandem pairs that encompass small RNA segments (186-399 nt) resembling Zetavirus-like sequences. *In vivo* characterization of the 5’-UTR with a ribozyme from a chrysovirid revealed that the RNA-cleaving activity is essential for protein translation initiation in fungi. Analogous experiments in plants with diverse ribozyme motifs indicated that just the presence of a self-cleaving activity can induce cap-independent translation. We conclude that RNA self-cleaving activity, historically linked to the rolling circle replication of viroid-like circRNAs, appears to be co-opted by linear RNA viruses for translational roles.

## Introduction

RNA’s dual nature as carrier of genetic information and biochemical catalyst has established it as a candidate molecule for the origin of life. RNA predates DNA, proteins and the genetic code in the primordial “RNA world” hypothesis^1–3^. Genetic remnants of this hypothetical RNA world would be the extant catalytic RNAs, such as the ribosomal RNAs, the RNAse P, or the small self-cleaving ribozymes. Therefore, the simplest RNA-based entities, such as viruses, viroids and other minimal mobile RNA genetic elements, are hypothesized to be living fossils of Earth’s first life^4,5^. The simplest known ribozymes belong to the family of small (50-200 nt) self-cleaving RNAs, with nine well characterized members: the hammerhead (hhrbz)^6,7^, hairpin (hprbz)^8^, human Hepatitis Delta (dvrbz)^9^, Varkud-Satellite (vsrbz)^10^, glmS^11^, twister (twrbz)^12^, twister sister, hatchet and pistol^13^ ribozymes. During the ‘80s, the first small ribozymes were described in infectious circRNAs of plants (hhrbz and hprbz in viroid-like agents), animals (dvrbz in Hepatitis Delta Virus) and fungi (vsrbz in a *Neurospora crassa* plasmid), but are also encoded in some DNA genomes^14,15^. In the last decade, these motifs have been discovered to be widespread in the DNA genomes of phages, bacteria, or eukaryotes^16–18^, including the human genome^19,20^. The precise biological functions of many of these genomic ribozymes are not well understood, but numerous hhrbz, dvrbz and twrbz motifs strongly associate with autonomous and non-autonomous retrotransposons across plant and metazoan genomes^21–26^.

Likely due to its simplicity^27^, the hhrbz is one of the most frequent ribozymes detected in nucleic acids. It is composed of three double helixes (I to III) that surround a core of 15 conserved nucleotides, and folds into a γ-shaped helical junction where the loops of helix I and II interact^28,29^. Depending on the open-ended helix, three main circularly permuted topologies (type I, II, or III) are possible for the hhrbz (Fig. 1A), although examples of novel permuted forms have also been recently reported for this ribozyme^30,31^. The dvrbz, another small ribozyme widespread among DNA genomes, shows a characteristic nested double pseudoknot structure with five helical regions (Fig. 1B).

**Fig. 1.**
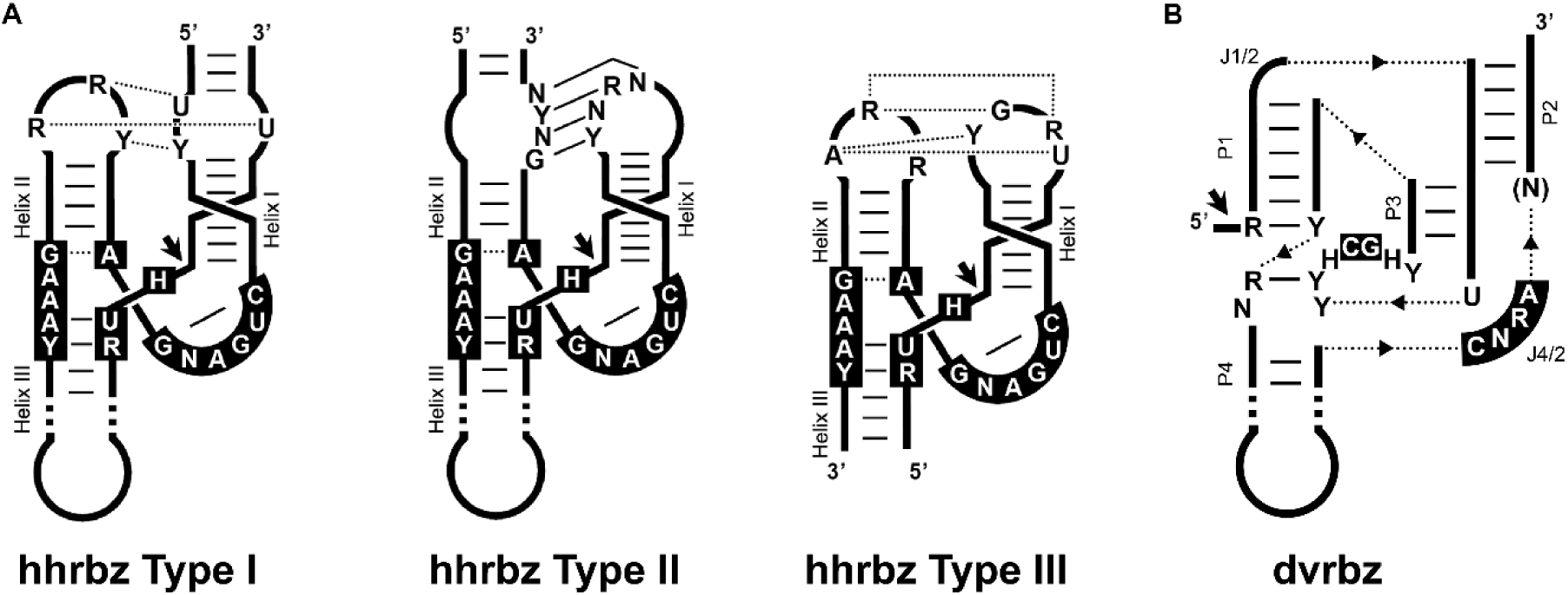
Schematic representation of (A) the three hammerhead topologies (hhrbz Type I, II, and III) and (B) the hepatitis delta virus (dvrbz) ribozymes. The conserved nucleotides are shown in black boxes. Typical tertiary loop-loop interactions of the hhrbz are indicated (dotted and continuous lines refer to non-canonical and Watson–Crick base pairs, respectively). Dotted lines with triangles represent connections between the helixes. Helical stems and single-stranded junction strands are indicated. Black arrows indicate the self-cleavage site.

Sequencing and computing have driven an exponential growth in the characterized biodiversity of microbial life forms. This expansion has relied on the use of protein hallmark genes or ribosomal RNA. However, we are starting to appreciate the existence of an extant and highly diverse RNA world of minimal agents. Previous analyses have unveiled hundreds of novel circRNA genomes with self-cleaving ribozymes in either one or both polarities^32,33^. More recently, these minimal circRNA agents were expanded to thousands of viroid-like taxonomic units, including examples of novel RNA virus-viroid hybrids in fungi^31,34^ or viroid-like obelisks in bacteria^35^. These discoveries blur the distinction between viroidal agents, ribozyme-bearing satellite viruses, and RNA viruses, but confirms the existence of a “modern RNA world” of mobile genetic elements of circRNA with and without small self-cleaving ribozymes^31^.

Here, we extend the conserved occurrence of small self-cleaving ribozymes, notably specific variants of the type I hhrbz, to diverse families of viruses with linear RNA genomes from fungi and plants. These catalytic motifs do not appear to be involved in the classical role of RNA processing of intermediates during rolling-circle replication, but instead have been functionally co-opted to perform novel translational roles in the life cycle of linear RNA viruses.

## Results

### Type I hammerhead ribozymes in dsRNA viruses

*In silico* screening for the nine known families of small ribozyme structures in viral sequences deposited in public databases revealed conserved motifs across evolutionarily diverse RNA viruses (see Methods for full details). Notably, putative ribozyme motifs were identified in the *Chrysoviridae* family of multipartite dsRNA viruses, where 42% (96 out of 226 contigs >2 kb) of the alphachrysovirus sequences contain a distinct variant of the type I hhrbz class (Fig. 1). These motifs are usually located in the 5’-UTRs (positive strand) of each of the four RNA segments comprising the viral genome (Fig. 2A and Supplementary Table 1). Exceptionally, a few alphachrysoviral segments exhibit a second hhrbz at the 3’-UTR region of the RNA (ie. Raphanus sativus chrysovirus 1, RNA segments 1 and 3. Supplementary Table 1). However, no examples of small ribozymes were detected in any of the betachrysovirus sequences analyzed (134 contigs >2 kb).

**Fig. 2.**
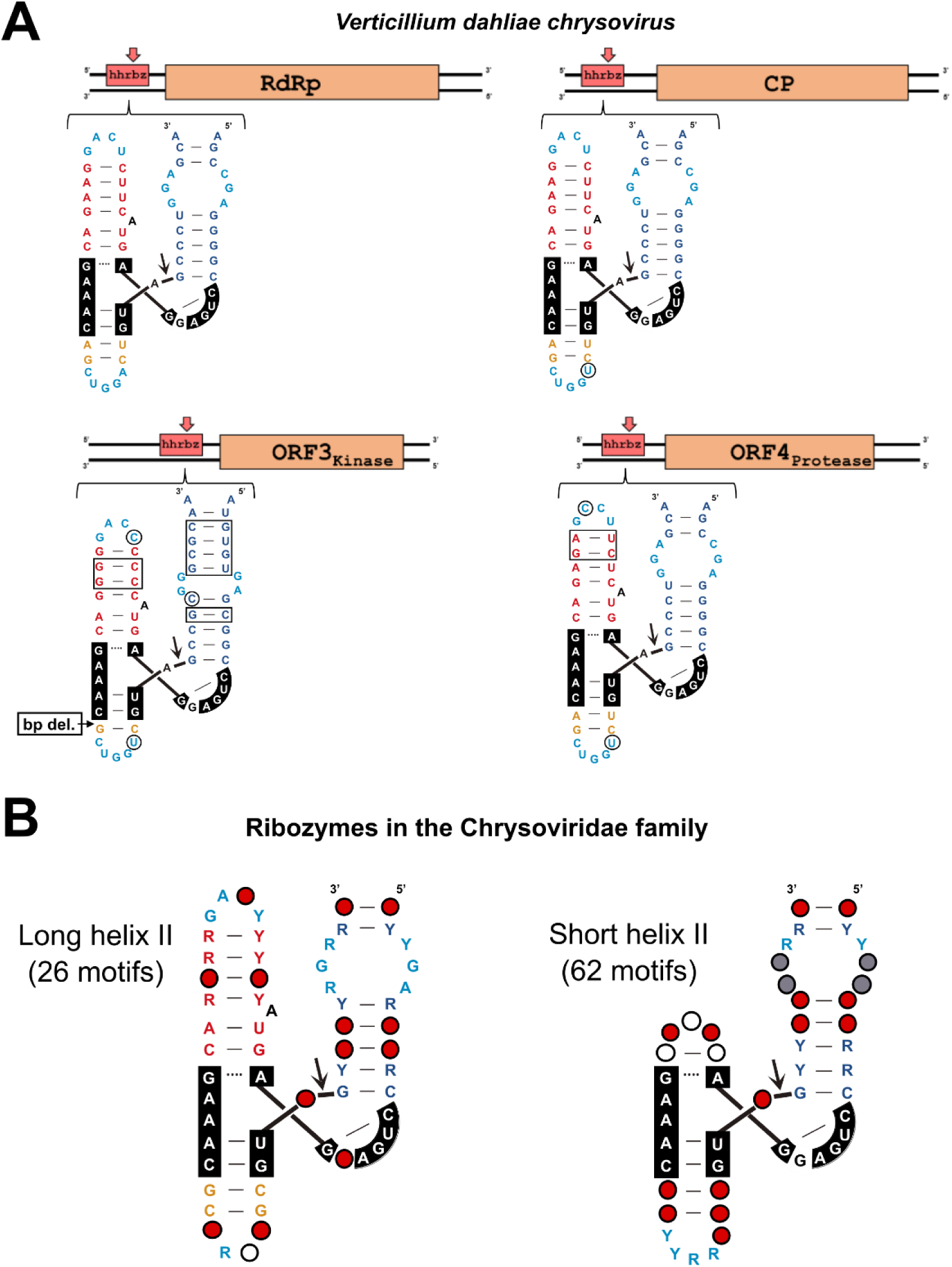
Fungal and plant chrysovirus hammerhead ribozymes (hhrbz). (A) The multipartite Verticillium dahliae chrysovirus^38^contains type I hhrbzs in the 5’-UTRs of the 4 RNA segments that make up the viral genome. Nucleotide changes in the ribozyme motifs with respect to the hhrbz at the RNA 1 encoding the RNA-dependent RNA polymerase (RdRp) are indicated by circles (single changes) and rectangles (base-pair covariations) (B) A summary of the covariance models for the two hhrbz architectures detected among most chrysoviruses. Red, grey and white dots refer to nucleotides present in more than 97%, 75% or 50% of the sequences, respectively. Y: C or U. R: G or A (See Supplementary Fig. 1A for details). CP, coat protein. ORF, open reading frame.

Most type I hhrbzs have been previously reported in diverse eukaryotic retroelements^14,15,22,25,36^, where they usually show a short helix III that prevents efficient self-cleavage of monomeric but not of dimeric hhrbz molecules^6,37^. However, chrysoviral type I hhrbzs show a longer helix III, resembling type I hhrbzs encoded in the DNA genomes of some bacteria/phages^16^, fungi^22^ and mammals^19^. The newly uncovered hhrbzs from chrysoviruses can be categorized into two conserved architectures with distinct properties (Fig. 2B); one third of the hhrbzs have long and stable helix II of 6 bp capped by a tetraloop and usually harboring a bulged adenosine in the middle of the stem, whereas two thirds show a short (1-2 bp stem) or even non-existent helix II (Fig. 2B and Supplementary Fig. 1).

*In vitro* analyses confirmed that chrysovirus hhrbzs with long helix II self-cleave efficiently during transcription (k_obs_∼2-3 min^−1^) (Fig. 3A). However, the observed rates of self-cleavage for the hhrbz with short (1 bp) or absent helix II were either low (k_obs_ ∼10^−2^ min^−1^) or very low (k_obs_ ∼10^−3^ min^−1^), respectively (Fig. 3B, C). These minimal rates of cleavage were observed during co-transcriptional experiments, but similar results were observed under post-transcriptional conditions, even at high pH (8.5) and Mg^+2^ concentrations (10 mM) (Supplementary Fig. 2).

**Fig. 3.**
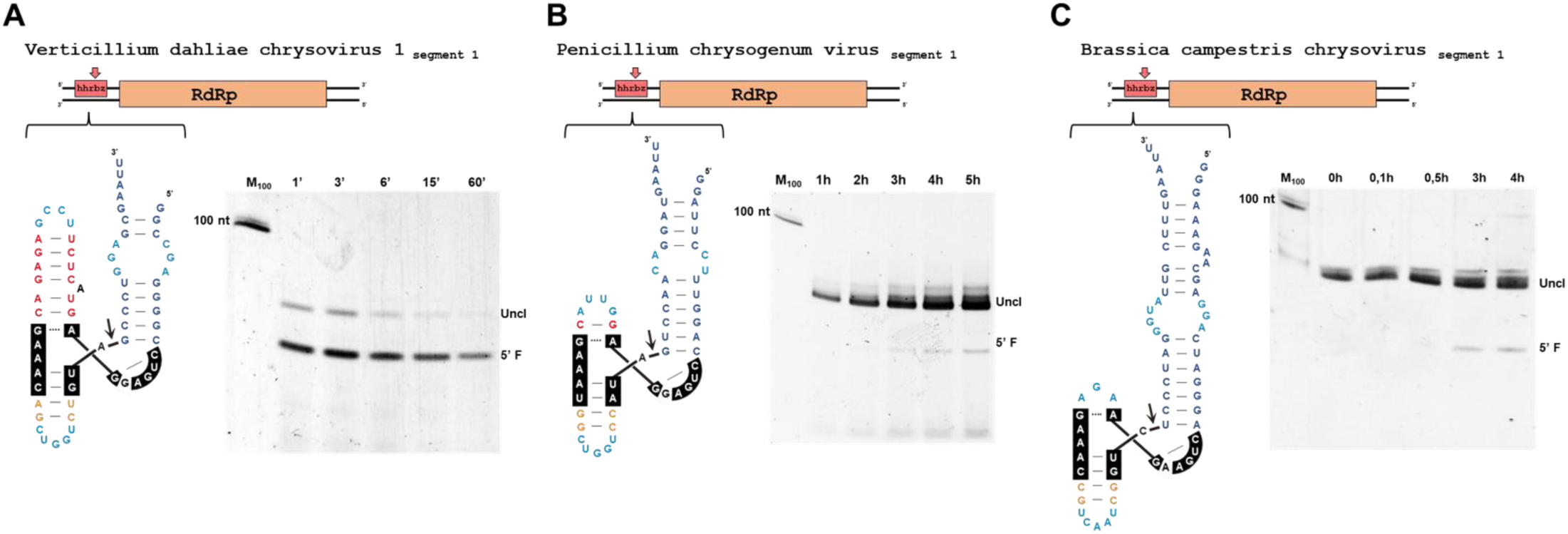
Self-cleavage kinetics of hammerhead ribozymes from fungal and plant chrysoviruses. (A) long helix II, (B) short helix II, or (C) totally absent helix II hammerhead ribozymes are catalytically competent *in vitro* but show distinct self-cleaving efficiencies. RdRp, RNA-dependent RNA polymerase. (A) and (B) correspond to co-transcriptional cleavage analysis under standard conditions (pH 7.5, 1 mM Mg^2+^), whereas (C) shows a post-transcriptional assay performed at higher pH (8.5) and Mg^2+^ concentration (10 mM) to increase its self-cleaving capability.

### Other ds and ssRNA viruses encode hammerhead, but also deltavirus and twister ribozyme motifs

Fusariviruses are a family of mycoviruses with monopartite ssRNA genomes that encode one to four ORFs^39,40^. Our bioinformatic analyses found that 114 of the 156 analyzed fusariviral sequences encode one or more type I hhrbzs, totaling 160 identified motifs (Supplementary Table 1). These hhrbz sequences and their helix sizes exhibit significant variability across viral genomes. As observed for chrysoviruses, they either follow a canonical type I hhrbz architecture, with medium size helixes I, II and III (4-8 bp stems), or variants with short helix II (1-2 bp stems) (Supplementary Fig. 1B, C). These observations suggest high and low self-cleavage efficiency, respectively, for each of the two architectures. Again, fusariviral ribozymes occur in the untranslated regions preceding most viral ORFs (Fig. 4A and Supplementary Fig. 3A). We made similar observations in the family of bipartite dsRNA megabirnaviruses^41^, which show the presence of slightly different variants of the type I hhrbz with longer helix II (Supplementary Fig. 2D, 3B). The motifs occur again in the large 5’-UTR of diverse megabirnavirus RNA segments (17 out of 32 analyzed contigs) (Supplementary Table 1). Interestingly, not only hhrbz motifs, but also an example of a *bonafide* dvrbz was detected in a genomic segment of the Rhizoctonia cerealis megabirnavirus-like RNA (Fig. 4B, Supplementary Fig. 2B). Similarly, instances of dvrbz, type I hhrbz and even twrbz motifs, were detected in the dsRNA genomes of diverse totiviruses. In these examples, however, the motifs were usually found as pairs of ribozymes (either from the same or different classes) arranged in close tandem separated by 186-399 nt (Fig. 4C). This arrangement echoes that observed in plant and animal retrozymes, where closely spaced tandem ribozyme pairs encoded in DNA genomes produce small circRNAs^24,25^. In the case of these totiviruses, the predicted circRNAs (i) can adopt stable rod-like structures, (ii) have nucleotide sizes that are multiples of three, and (iii) contain ORFs without stop codons, potentially encoding a never-ending polypeptide in the positive strand (Fig. 4C, Supplementary Fig. 4). These three features somewhat resemble those described for the intriguing group of viroid-like Zetaviruses genomes detected in diverse environmental metatranscriptomic studies^31,32^ (Supplementary Fig. 4D). However, predicted circRNAs in totivirus-like genomes usually lack a key characteristic of Zetaviruses, as they do not retain any of the flanking ribozymes within their circular sequences^31,34^.

**Fig. 4.**
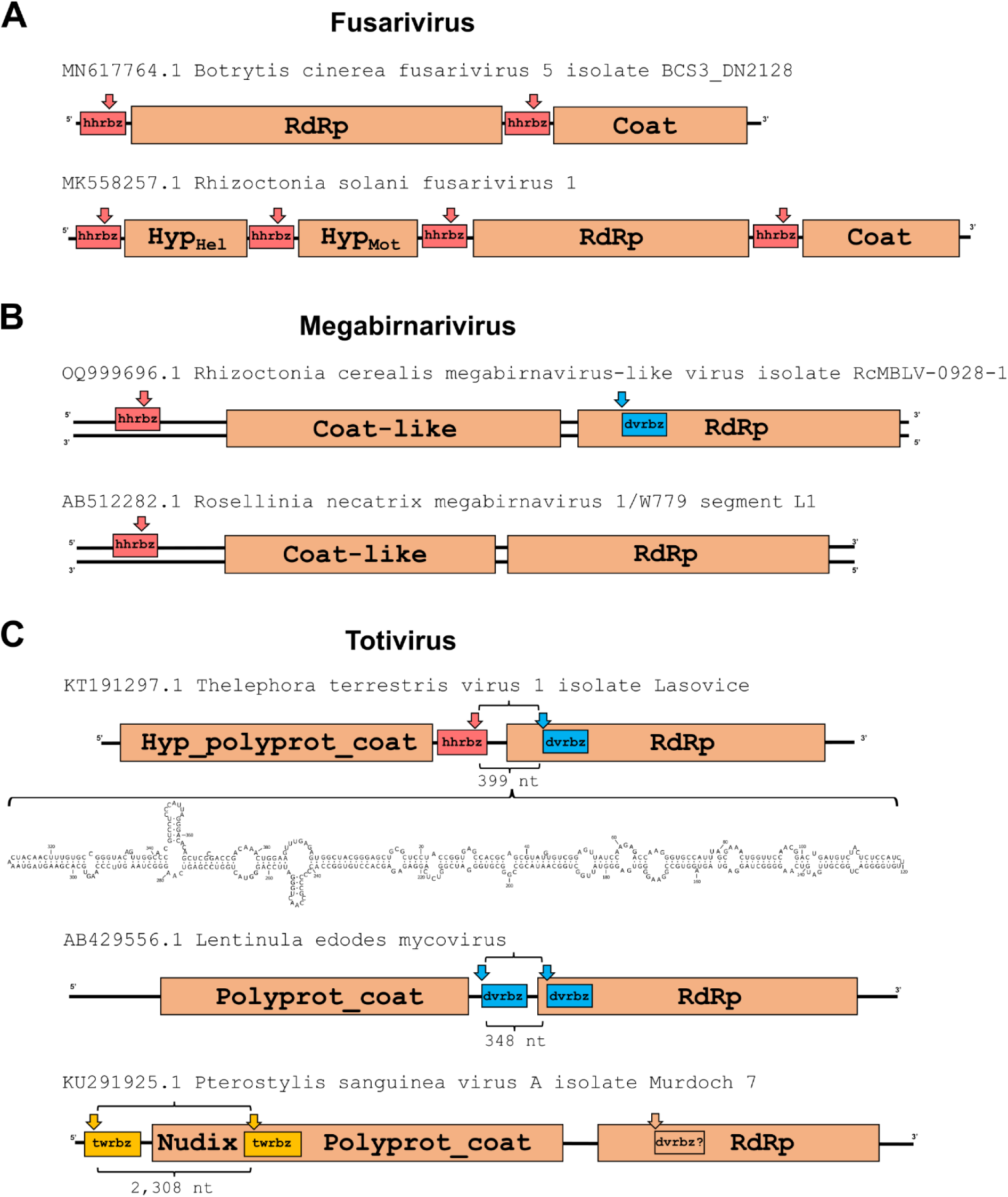
Small ribozymes are widespread in fungal ss and dsRNA viruses with linear genomes. (A) fusariviruses, (B) megabirnaviruses and (C) totiviruses encode conserved self-cleaving ribozymes of the hhrbz, dvrbz and twrbz classes. In totivirus genomes, pairs of hammerhead and delta self-cleaving ribozymes occur close to each other (∼189-399 nt). Similar arrangements of close ribozymes in tandem responsible of circRNA formation have been previously identified in plant^24^ and metazoan^25^ genomes, suggesting that totivirus ribozymes would allow the generation of small circRNAs, as depicted for Telephora terrestris virus 1 (See Supplementary Fig. 4 for more details).

Our bioinformatic analyses also revealed the presence of analogous type I hhrbzs in the UTRs of some dsRNA genomes from additional fungal viral families, such as fusagraviruses and hypoviruses (Supplementary Table 1). Moreover, we also detected diverse variants of the twister ribozyme at the 5’-UTRs of some phlegi- (ie. Rhizoctonia zeae dsRNA virus 1 and 2)^42^ or tombus-like viruses from diverse soil and plant-associated microbial communities (Supplementary Fig. 5). Altogether, these results confirm the prevalence of small self-cleaving ribozymes encoded within the linear RNA genomes of diverse fungal and plant viruses.

### Hammerhead ribozymes in RNA viruses show self-cleaving activity *in vivo*

Given the widespread occurrence of type I hhrbzs among chrysovirids and fusarivirids, we chose to look for *in vivo* evidence of ribozyme self-cleaving activity during viral infection. We selected two chrysoviruses (either infecting a plant or a fungus) and one fungal fusarivirus as examples of ds and ssRNA viruses with hhrbzs in different 5’-UTRs regions, respectively.

Brassica campestris chrysovirus 1 (BcCV1) is a tripartite dsRNA chrysovirus infecting brassica plants^43^, which shows conserved hhrbzs (short helix II variants with a minimal self-cleaving activity *in vitro*, see above) in the 5’-UTRs of each RNA segment. The expected cleavage sites of the hhrbzs map at positions 58, 60 and 60 of RNA segments 1, 2 and 3, respectively (Supplementary Fig. 6A). Rapid amplification of cDNA ends (RACE) experiments using total RNA from a BcCV1-infected *B. oleracea* plant generated two amplification products for each of the chrysovirus RNAs, which are more evident in the case of RNA2 and RNA3 (Supplementary Fig. 6B). Cloning and sequencing of the amplicons confirmed the existence, for each genomic RNA segment, of molecules with two different 5’ ends, one corresponding to the expected viral full-length RNA segment, and the other one with the self-cleavage site predicted for the respective embedded ribozyme (Supplementary Fig. 6C).

The recently characterized Gnomognopsis castanea chrysovirus 1 (GcCV1) has 4 genomic segments encoding one ORF each, except for the bicistronic RNA3^44^. We detected the presence of 5 different hhrbz motifs (long helix II variants with efficient self-cleaving activity *in vitro*, see above), one in each 5’-UTR preceding an ORF (Supplementary Fig. 7). Here, RACE experiments carried out on total RNA extracted from an infected fungus isolate showed that the population of viral RNAs is also composed of RNA segments with two different 5’ termini, as supported by clones either mapping to the expected 5’ end of the genomic RNA, or to the predicted 5’ site of self-cleavage of each hhrbz (Supplementary Fig. 7). Such a mix of RNA molecules with two different ends is particularly evident for RNA2, which gives two abundant and distinct bands in PCR RACE amplifications (Supplementary Fig. 7E). It is of note that, in this virus, the two copies of hhrbz present in each one of the two UTRs of the bicistronic RNA3 were found to self-cleave *in vivo* at their predicted sites.

Finally, Pleospora tiphycola fusarivirus 1 (PtFV1) is a ssRNA virus present in an ascomycetes fungus isolated from the seagrass *Posidonia oceanica*^45^. Members of the recently established *Fusariviridae* family are a group of non-segmented viruses with heterogeneous genome organization according to the genus they belong to^40^. The RNA genome of PtFV1 can potentially express up to three proteins from three distinct ORFs^45^. It also contains a predicted type I hhrbz with a short helix II, similar to the Penicilium chrysogenum virus ribozyme (Fig. 3B). The catalytic RNA essentially spans the entire small intergenic UTR (66 nt), with the self-cleavage site (position 4,735) just 7 nucleotides upstream of the proposed ORF2 start codon (position 4,742). RACE analysis using reverse transcription products primed around 200 nt downstream the cutting site confirmed the existence *in vivo* of an RNA species with the predicted 5’ end after self-cleavage (Supplementary Fig. 8A). Moreover, northern blot analysis showed the accumulation of a positive-sense RNA fragment of a size compatible with that of a subgenomic RNA originated by self-cleavage of the ribozyme (Supplementary Fig. 8B). Overall, experiments with total RNA purified from either infected plants or fungi followed by RACE analyses confirm that the population of viral RNA genomes *in vivo* is heterogeneous, being composed of uncleaved and self-cleaved RNA molecules.

### Hammerhead ribozyme self-cleavage in a chrysovirus UTR allows cap-independent translation

Translation initiation activity, presumably via some kind of internal ribosome entry site (IRES) element, has been proposed for the 5’-UTR and subsequent 72 nt of coding region of fungal dsRNA viruses such as chrysoviruses and hypoviruses^46^. Among these, the 5’-UTR of the alphachrysovirus Helminthosporium victoriae virus 145S (HvV145S) RNA2 was suggested to carry an IRES-like element. We found that, as in most alphachrysoviruses, the 5’-UTRs of the HvV145S RNA segments also contain conserved type I hhrbzs (Supplementary Table 1, Fig. 5A). These ribozymes correspond to the hhrbz variants with long helix II (Fig. 2B), which are expected to reach efficient self-cleaving activity *in vitro* and *in vivo*. To understand whether hhrbzs are directly involved in protein translation initiation, on- and off-target mutations were introduced into the self-cleaving motif and analyzed *in vivo*. The transgenic expression of bicistronic RNAs in *Cryphonectria parasitica* mycelia was carried out as previously described^46^, in which the upstream ORluc and downstream OFluc genes were translated in a cap-dependent and cap-independent manner, respectively (Fig. 5B). The original HvV145S-2 5’-UTR sequence and its truncated variants in the 5’ proximal region preceding the ribozyme (Δ60 and Δ120) showed similar ratios of OFluc to ORluc chemiluminescence intensity (RLU), suggesting that this off-target 5’ region is dispensable for translation initiation (Fig. 5D). In contrast, the deletion of the whole region corresponding to the hhrbz motif (Δ124-183) abolished translation initiation of OFluc, resulting in a significantly lower OFluc/ORluc RLU ratio. In addition, either a single point mutation in the UTR ribozyme at position G159, the general base catalyst in the self-cleavage reaction^29^, or deletion of 5 essential nucleotides, including the conserved U-turn motif^47^ of the catalytic core of the ribozyme (Δ134-138), supressed the translation initiation ability observed for the wt 5’-UTR sequence carrying the catalytically competent hhrbz (Fig. 5C and D).

**Fig. 5.**
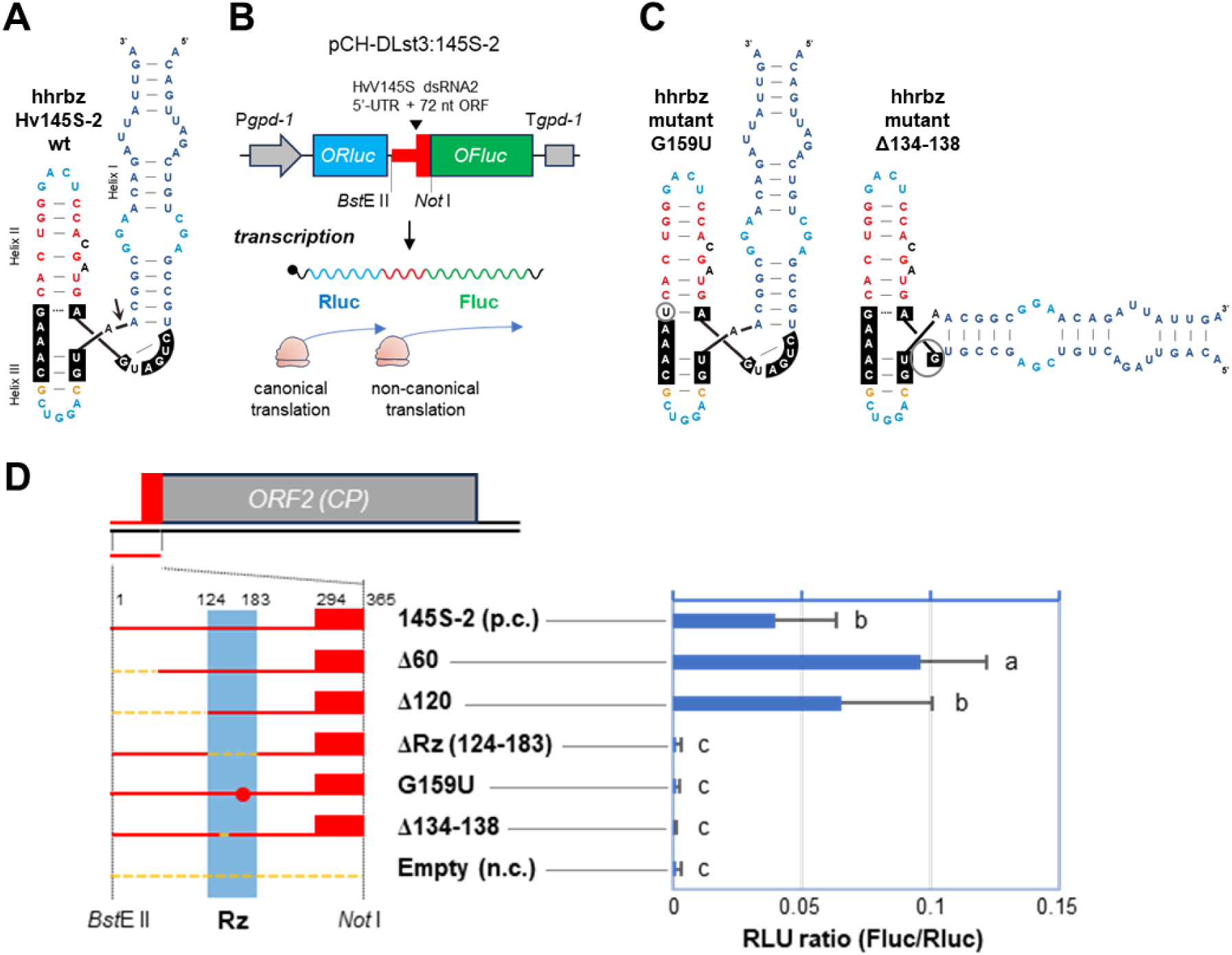
Mutational analyses of the hammerhead ribozyme present in the 5’-UTR of the chrysovirus HvV145S RNA2. (A) Predicted structure of the HvV145S hammerhead ribozyme (hhrbz) present in the 5’-UTR of the RNA2. (B) A bicistronic dual-luciferase reporter system in fungal mycelia for investigation of the cap-independent initiation activity located at the 5’-UTR and coding region (72 nt) of HvV145S dsRNA2. Codon-optimized Renilla luciferase (ORluc, blue box) and firefly luciferase (OFluc, green box) genes are translated either in a canonical (cap-dependent) or a non-canonical (cap-independent) manner, respectively. Red line and box indicate HvV145S-2 sequence. (C) Predicted secondary structures of the 145S-2 hhrbz mutants containing either a substitution (left, G159U) or a deletion (right, Δ134-138), as indicated with grey circles. (D) Left, schematic representation of the 5’-UTR and first 72 nt of coding region (in red) of HvV145S-2 (in grey) used for the bicistronic dual-luciferase reporter system. The details of the mutations are represented at the bottom as the sequence variant names. Right, dual luciferase reporter assay results for each construct showing the ratio of luminescent intensities (Fluc RLU/Rluc RLU). p.c., positive control, n.c., negative control.

### Diverse small self-cleaving RNA motifs induce translation initiation in plant cells

To further investigate whether the observed effect of the chrysoviral hhrbz on protein translation is specifically due the RNA self-cleaving activity or any other domains present within the viral UTR, we employed a similar dual-luciferase reporter system designed for its expression in agroinfiltrated *N. benthamiana* plant cells^48^. The bicistronic constructs contained the Rluc and Fluc genes under the control of 35S promoter and separated by a small (∼150 bp) non-coding spacer either containing a standard Multiple Cloning Site sequence (85 bp) or different small self-cleaving motifs (∼85-110 bp), such as type I hhrbzs from a plant (BrcCV1) and a fungal (VdCV1) chrysovirus, from a megabirnavirus (RnMBV1) or from a human gene intron (the HH9 motif^19^). In addition, a dvrbz from an animal deltavirus (NewtDV, genomic sequence) was also assayed (Fig. 6A). As expected, the construct carrying the MCS sequence chosen as a negative control showed no noticeable OFluc/ORluc ratio (Fig. 6B). However, constructs with chrysoviral hhrbz motifs produced a weak but discernible signal, while a much higher ratio was observed with the Rosellinia necatrix megabirnavirus 1 (RnMBV1) hhrbz. Moreover, a construct carrying the dvrbz also showed a significant signal of OFluc protein expression (∼0.4%). In contrast, assays using the HH9 ribozyme, a fast self-cleaving hhrbz^19^, resulted in minimal Fluc/Rluc ratios. However, it should be noted that both Fluc and Rluc raw signals for the construct containing the HH9 ribozyme were consistently one order of magnitude lower than the observed one for other constructs, suggesting that HH9 activity interferes with the normal expression of the reporter system.

**Fig. 6.**
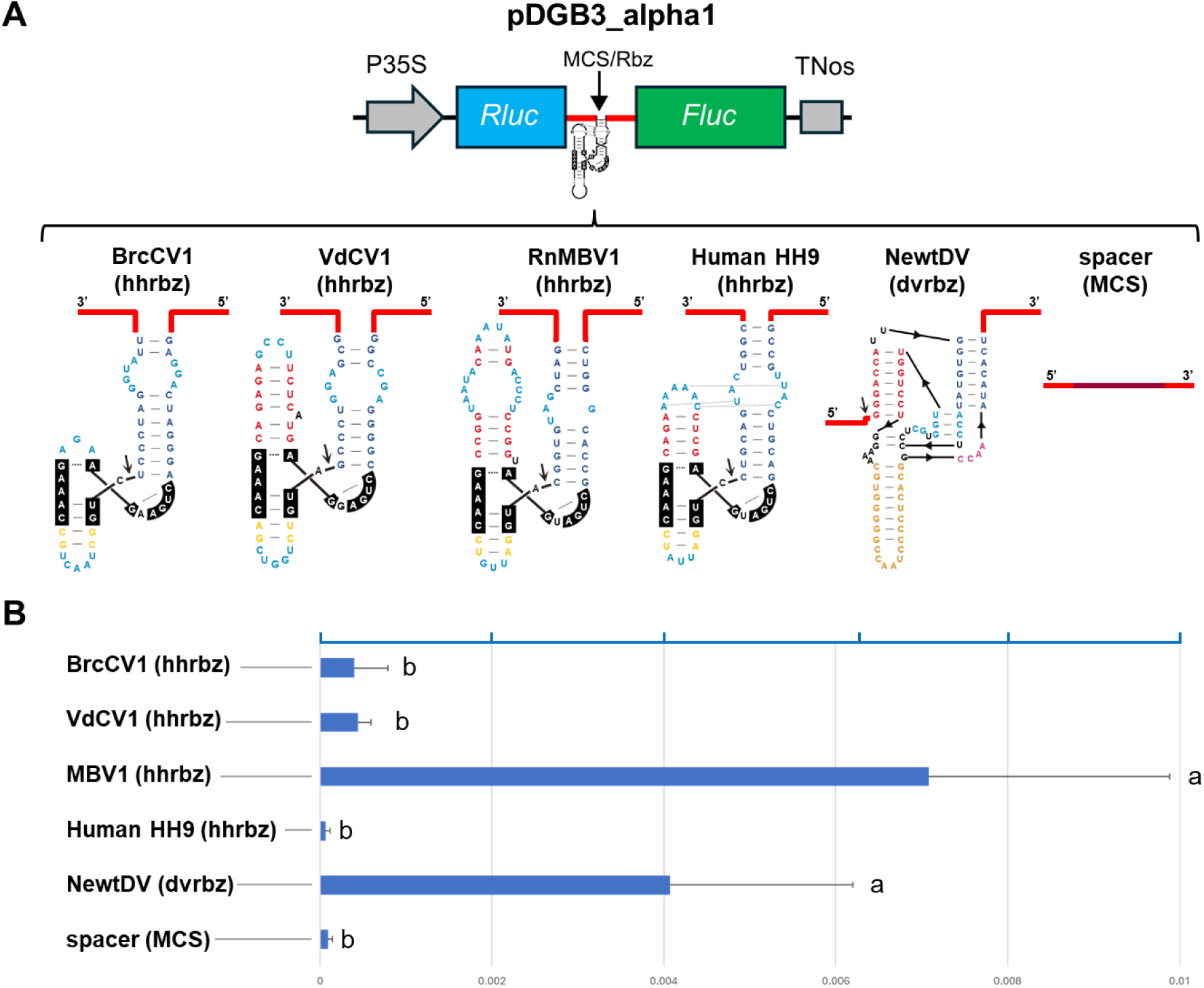
Analysis of RNA self-cleavage on a dual-luciferase reporter expression in *N. benthamiana* leaves. (A) Schematic representation of the dual-luciferase reporter construct used in the study, with the Rluc and Fluc genes separated by a short DNA region (∼130 bp) including either a MCS sequence (49 bp) or various self-cleaving RNA motifs, such as hammerhead ribozymes (hhrbz) from plant (BrcCV1) and fungal (VdCV1) chrysoviruses, a megabirnavirus (RnMBV1), a human gene intron (HH9), a deltavirus ribozyme (dvrbz) from NewtDV, and a negative control construct just containing MCS sequence. (B). dual luciferase reporter assay results for each construct agroinfiltrated in plants showing the ratio of luminescent intensities (Fluc RLU/Rluc RLU).

## Discussion

Almost four decades after their discovery^6,7^, the origins and evolutionary history of small self-cleaving ribozymes remain a complex puzzle. It is assumed that tiny RNAs capable of sequence-specific cleavage and ligation could be among the first catalytic motifs to emerge during the RNA world, with a role in the replication of primordial RNA genomes^49^. Our knowledge about small ribozymes has been mostly limited to those from a unique family of a few viroid-like circRNAs (∼15 species). In the last years, however, small catalytic RNAs have been found as ubiquitous motifs encoded in the DNA genomes of organisms throughout the tree of life^16,50^ and, more recently, within thousands of novel viroid-like circRNAs in either eukaryotic or prokaryotic hosts^31–35^. Thus, this plethora of catalytic motifs are likely expected to have more biological roles beyond multimer self-processing.

In this work, we have expanded our knowledge about the occurrence of small ribozymes to typical ss- and dsRNA viruses with linear genomes. Notably, hundreds of novel variants of the type I hhrbz are found to be conserved in the 5’-UTRs of fungal and plant RNA viruses belonging to the *Chrysoviridae* and *Fusariviridae* families. It should be highlighted that nearly all hhrbz motifs previously detected in infectious circRNAs, such as plant viral satellites and viroids^51^, obelisks^35^, deltaviruses^52^, ambiviruses or mitoviruses^31,34^, correspond to the type III architecture, whereas type I hhrbzs are usually DNA-encoded motifs present in the genomes of either bacteria and bacteriophages^16,17,53^ or metazoans^19,22,25,36^. On the other hand, natural hhrbzs adopt a similar three-way junction fold with conserved loop-loop interactions between helixes I and II (Fig. 1), which are crucial to achieve high activity under *in vivo* low magnesium concentrations ^28,29,54^. The new viral type I hhrbzs described in this work, however, do not seem to have kept any of the typical loop-loop interactions conserved in known hhrbzs (Fig. 1) or in other three-helical RNAs with remote tertiary contacts^55^. Moreover, stem lengths of helix II in viral hhrbzs are either slightly longer (6-8 bp) or shorter (0-2 bp) than the ones present in most type I hhrbzs (4-5 bp) (Fig. 2B). These observations, together with a medium to low *in vitro* self-cleaving activity for viral hhrbzs, point to a lack of any clear loop-loop interactions in these motifs. These viral hhrbzs are however functional *in vivo*, as revealed by the co-existence of uncleaved and cleaved RNA genomes in plant and fungal infected cells, indicating that viral sub-genomic RNAs arise through a novel mechanism catalyzed by small self-cleaving ribozymes.

Regarding their biological role, most of the hhrbzs reported here are small motifs (50-70 nt), which nevertheless can span a significant portion of the viral UTR sequences (usually ∼100-300 nt) where they are embedded. It has been proposed that, among other functions, these 5’-UTR are involved in cap-independent translation initiation acting as novel IRES-like elements^46^. Our deletion mutants of a chrysoviral UTR show that the hhrbz motif (nucleotides 124-182) is specifically required for translation initiation function. More intriguingly, a single point mutation in the catalytic G base within the hhrbz core (known to inhibit RNA catalysis without altering the whole ribozyme structure^56^) was enough to abolish the protein expression observed with the wild-type UTR. This result suggests that self-cleaving activity would be crucial for the role of the UTR in cap-independent translation. Consequently, we advance that viral ribozymes may allow translation initiation thanks to their self-cleaving activity through an unknown mechanism. As a second alternative, we can also envisage that a mutation preventing self-cleavage may indeed affect, albeit minimally, the fold of the hhrbz, which could disrupt a potential role as an IRES-like structure. To better understand the role of RNA catalysis in eukaryotic translation initiation, we moved to a similar bicistronic reporter system using plants instead of fungal protoplasts. This new approach was based solely on the presence of isolated small ribozyme motifs within dual-luciferase constructs and in the absence of any additional viral UTR sequence. Interestingly, the presence of viral self-cleaving motifs alone was, in most cases, sufficient to promote detectable translation initiation in plants when compared with the construct lacking any ribozyme. These results, however, should be interpreted with caution. First, the translation ratios of classic luciferase genes obtained in plants with ribozyme-only constructs were modest (usually below 1%) relative to those achieved with the full chrysoviral UTR (including 72-nt coding region) obtained with codon optimized luciferase genes in fungi^46^ (∼4%-8%). Second, not all ribozymes behaved equally, as translation levels were really low for the chrysovirus hhrbz (∼0.05%) but more evident for the megabirnavirus hhrbz or the newt dvrbz (0.4– 0.5%). Third, the hhrbz derived from the human genome (HH9) not only failed to enhance Fluc expression but caused an overall reduction in luciferase activity from both reporters, suggesting a detrimental effect on the whole mRNA stability and expression. Altogether, these results indicate that the self-cleaving activity can promote translation initiation, although the magnitude of this effect appears to depend on the specific ribozyme context, biological system and self-cleaving efficiency. In this regard, direct links between RNA self-cleaving motifs and eukaryotic protein translation have been already advanced in the literature. Notably, the dvrbz present in the 5’ end of various eukaryotic retrotransposons has been reported to promote protein translation initiation^21^. Consequently, this activity would mirror the mechanism we describe here in RNA viruses.

The noticeable differences in protein expression found for different catalytic RNA motifs suggest that self-cleaving activity could be a necessary condition but not sufficient. These observations imply that the precise ribozyme sequence and/or structure may jointly contribute to translation initiation. In this context, the global architecture and topology of the type I hhrbz detected in RNA viruses shows clear structural similarity with the core domains of diverse viral IRES elements, such as those from picornaviruses^57^, the encephalomyocarditis virus (EMCV)^58,59^ or the Hepatitis A virus (HAV)^60,61^ (Supplementary Fig. 9), suggesting that not only the self-cleaving activity but also the hhrbz ribozyme fold itself may have a potential role in the recruitment of some translation initiation factors. This possibility could be extended to the dvrbz, which despite clear structural differences with the hhrbz, they all share a similar compact three-helical junction topology (where two helices coaxially stack and the third helix runs parallel to the major axis of the Y-like structure) characteristic of the core domain of diverse IRESs (Supplementary Fig. 10)^55^.

Altogether, our data reveal a novel and unexpected activity in linear RNA viruses, whereby the genomic RNA undergoes self-cleavage at one or more sites during the viral life cycle. This cleavage sacrifices full genomes to generate sub-genomic RNA molecules no longer suitable for further replication, making a controlled balance between cleaved and intact genomes crucial to preserve enough replication templates.

In this regard, most ribozymes identified here are far from optimal motifs (ie. type I hhrbz seem to lack crucial tertiary interactions^28^), resulting in medium to low self-cleaving activity *in vitro*. Such deviation may allow a significant fraction of viral RNA to evade cleavage and remain competent for replication. Meanwhile, sub-genomic RNA fragments resulting from self-cleavage would serve as templates for translation initiation through an unclear cap-independent mechanism. Although further research will be needed to understand this mechanism, our findings shift the traditional paradigm of small ribozymes as mere processing elements for rolling circle replication of circRNA agents, expanding the evolutionary toolkit of RNA viruses and their biotechnological potential.

## Methods

### Bioinformatic analysis of viral sequences

Computational searches were performed with the INFERNAL 1.1 software^62^ using covariance models representing known small-ribozyme classes, and their topological variants, that were obtained from the RFAM 14 database^63^. In addition, newly developed covariance models were included to account for recently identified ribozyme architectures recently described^30,31^. A first screening of a collection of 852 thousand contigs (with lengths ranging between 2,000 and 50,000 bp) of DNA or RNA viral origin retrieved from the Genbank NR database (2024) resulted in hundreds of ribozyme hits that were manually inspected. Ribozyme hits present in viral vectors, patented sequences, DNA bacteriophages^16^ or in RNA ambiviruses and mitoviruses^31^ already described in circRNA replication were discarded, whereas those hits with low E-values such as type I hammerhead (E-val < 10^−3^), deltavirus (E-val < 0.1) or twister (E-val < 10^−3^) ribozymes detected in RNA viruses from diverse families were used to build new covariance models to carry out re-screenings (https://github.com/delapenya/RNA_virus_rbz). Analyses of specific RNA viral families showing the presence of small ribozymes and refinement of covariance models were performed iteratively.

The consensus features of the ribozyme motifs were calculated based on the ribozyme sequences obtained for each family. Nucleotide frequencies and covariation, as well as the corresponding drawing, were the output of the R2R software^64^. Multiple sequence alignments were performed using MUSCLE^65^.

### Cloning of ribozyme sequences and transcriptions

The sequences corresponding to the hhrbz motifs of the RNA 1 segments of Verticillium dahliae chrysovirus 1 (VdCV1), Penicilium chrysogenum virus and Brassica campestris chrysovirus 1 (Fig. 3) preceded by the T7 RNA polymerase promoter were purchased as gBlock Gene Fragments (Integrated DNA Technologies). The DNA fragments were cloned into a linearized pUC18 vector by BamHI and EcoRI cohesive-end ligation. RNAs of the cloned sequences were synthesized by *in vitro* run-off transcription of the linearized plasmids containing the hhrbzs. The transcription reactions contained 40 mM Tris–HCl, pH 8, 6 mM MgCl_2_, 2 mM spermidine, 0.5 mg/ml RNase-free bovine serum albumin, 0.1% Triton X-100, 10 mM dithiothreitol, 1 mM each of ATP, CTP, GTP, and UTP, 2 U/µl of Ribonuclease Inhibitor (Takara Inc), 20 ng/µl of plasmid DNA, and 4 U/µl of T7 RNA polymerase. After incubation at 37°C during the indicated time, the products were fractionated by 10% polyacrylamide gel electrophoresis (PAGE) in the presence of 8 M urea.

### Kinetics of self-cleavage

Analyses of hhrbz self-cleaving activity under co-transcriptional and post-transcriptional conditions were performed as previously described^66^. Appropriate aliquots of the transcription reactions (progressive smaller volumes were taken at longer incubation times in co-transcriptional assays) were removed at different time intervals, quenched with a fivefold excess of stop solution (8 M urea, 50% formamide, 50 mM EDTA, 0.1% xylene cyanol and bromophenol blue) at 0°C, and analyzed as previously described^22,52^. For co-transcriptional cleavage analysis, the uncleaved and cleaved transcripts obtained at different times were separated by PAGE in 10% denaturing gels and detected by Sybr Gold staining (Thermo Fisher Scientific). For post-transcriptional cleavage analysis, uncleaved primary transcripts were eluted by crushing the gel pieces and extracting them with phenol saturated with buffer (Tris–HCl 10 mM, pH 7.5, ethylenediaminetetraacetic acid [EDTA] 1 mM, sodium dodecyl sulfate 0.1%), recovered by ethanol precipitation, and resuspended in deionized sterile water. To determine the cleaving rate constants, uncleaved primary transcripts (from 1 nM to 1 µM) were incubated in 20 µl of 50 mM Tris–HCl at the appropriate pH (either 7,5 or 8,5) for 1 min at 95 °C and slowly cooled to 25 °C for 15 min. After taking a zero-time aliquot, self-cleavage reactions were triggered by adding MgCl_2_ at a final concentration of either 1 or 10 mM. Aliquots were removed at the appropriate time intervals and quenched with a fivefold excess of stop solution at 0 °C. The substrate and self-cleaved products were separated by PAGE in 10% denaturing gels and detected by Sybr Gold staining (Thermo Fisher Scientific). In both cases, the product fraction at different times, Ft, was determined by quantitative scanning of the corresponding gel bands and fitted to the equation F_t_=F_∞_(1 − e^−kt^), where F_∞_ is the product fraction at the reaction endpoint, t is the time in minutes, and k is the first-order rate constant of cleavage (k_obs_).

### RACE and northern blot analyses

Analysis of fusarivirus-infected fungi was carried out using the isolate MUT4379 from the Mycotheca Universitatis Taurinensis. Gnomognopsis castanea chrysovirus 1 infected isolate 1 was recently described^44^. A symptomless plant of cabbage “Torzella” (*Brassica oleracea* L. var. acephala) infected with BcCV1 (isolate TC-3) was used for plant chysovirus 5’ Rapid amplification of cDNA ends (5’ RACE) analysis^67^.

Plant and fungal total RNA was extracted from leaves of cabbage “Torzella” plants and 4-day old mycelial liquid cultures grown in PDB following protocols previously described^44,67^. Hirtzman’s protocol for 5’ RACE was performed as previously described in detail^68^. For each RACE two different primers were designed downstream the predicted ribozyme cleavage site. Regarding GcCV1, the primers used to demonstrate the *in vivo* activity of the ribozymes found in the 5’ end of each genomic segment are reported in a previous work^44^. For the RACE demonstrating the *in vivo* activity of the ribozyme found in the intergenic region of the RNA3 of GcCV1, two RACE primers were designed (Supplementary Table 2); the oligonucleotides used to prime RACE for PtFV1 and BcCV1 are also displayed in Supplementary Table 2. Briefly, RACE analysis was performed on the 5′ ends of each viral sequence to confirm the presence of a self-cleavage site. The RACE method used^68,69^ rely on the synthesis of a cDNA using the specific primers mentioned before and the Superscript IV (Thermo Fisher Scientific, Waltham, MA, USA) reverse transcriptase. Resulting cDNAs were tagged with polyG or polyA using deoxynucleotidyl transferase (Promega, Madison, WI, USA). PCRs were then performed using primer complementary to the polyG or polyA added, and a second specific primer on the viral sequence (Supplementary Table 2). Obtained PCR bands were purified from electrophoresis gel using Zymo gel DNA recovery kit (Zymo Research, Irvine, CA, USA), inserted in a plasmid using the pGEMT easy vector kit (Promega, Madison, WI, USA). Transformed colonies were selected and obtained plasmids were purified using ZR Plasmid Miniprep kit (Zymo Research, Irvine, CA, USA) and sent for Sanger sequencing. For each RACE a minimum of 5 clones were sequenced.

Northern blot of fungal extracts for MUT4379 was performed as previously described^70^ using a riboprobe (detecting the positive sense viral RNA) obtained by *in vitro* transcription of a cDNA fragment cloned into pGEM-T-Easy (PROMEGA, Madison, WI) corresponding to the genomic region between nt 4,800 and nt 4,980 of the deposited fusarivirus sequence (NCBI NC_028470).

### Dual-luciferase (DL) reporter assays

The DL assay using fungal mycelia was performed as previously described^46^. Briefly, the pCH-DLst3 vector and PCR-amplified fragments of 5’-UTR and adjacent 72 nt of coding sequence (5’-UTR+) from HvV145S dsRNA2 were digested with *BstE* II and *Not* I, and ligated. The clone obtained was used for transformation of *Cryphonectria parasitica* EP155, and mycelia of transformants were subjected to the DL assay as described^46^. The vector pCH-DLst3 carries a codon-optimized Renilla (ORluc) and a Firefly (OFluc) luciferase genes in a bicistronic manner, under the control of the gpd-1 promoter/terminator system. *BstE* II and *Not* I recognition sites in between ORluc and OFluc were used for inserting the 5’-UTR and adjacent 72 nt coding cDNA sequence (5’-UTR+; original and mutant variants), expecting that ORluc and OFluc are translated in cap-dependent and cap-independent manner, respectively. *C. parasitica* EP155 was transformed with the different constructs and mycelia of transformants were subjected to DL assay using a dual-luciferase reporter assay system (Promega). A small patch of mycelia was homogenized in a microtube with 200 μl of Cell-Lysate solution, and 10 μl of solution were mixed with 100 μl of the firefly luciferase substrate solution, then the first luminescence intensity was measured using GloMax 20/20 (Promega) with a detection time of 1 s. This was followed by the addition of 100 μl of the Renilla luciferase substrate solution, and the second luminescence measurement. Translational activities were calculated as a ratio of OFluc RLU (relative luminescent unit)/ ORluc RL.

The approach used in fungal mycelia was adapted to *Nicotiana benthamiana* plants as previously described^48^. Briefly, a bicistronic construct was made with the two luciferase genes, Rluc and Fluc^71^, separated by three stop codons and a small region of ∼150 bp either containing a non-coding spacer (pUC Multiple Cloning Site, or MCS of 84 bp) sequence as a negative control, or different small self-cleaving motifs (∼85-110 bp). The entire construct was placed under the control of the 35S promoter and the Nos terminator^48^, and cloned into the binary vector pDGB3-alpha1_[NO_PRINTED_FORM]_ through 25 cycles of restriction and ligation reactions, using the restriction enzyme BsaI and T4 DNA ligase (Promega) .Each final plasmid was transferred into *Agrobacterium tumefaciens* strain C58 by electroporation^72^. Overnight-grown bacterial cultures were pelleted and resuspended in agroinfiltration medium, containing 10mM MgCl_2_, 10mM MES and 150µM acetosyringone, to an optical density at 600 nm of 0.1 to 0.2. For transient expression experiments, five-week-old *N. benthamiana* plants were inoculated and maintained under controlled conditions for 6 days. Extracts from 1 cm² of agroinfiltrated leaf tissue were prepared 5- and 6-days post-infiltration, using 400 μl of 0.1M Tris-HCl buffer (pH 7.8). Subsequently, 50 μl of extract were mixed with 45 μl of extraction buffer and 5 μl of 10mM luciferase substrate solution (coelenterazine or D-Luciferin to measure Rluc or Fluc activity, respectively). Stock solutions of the substrates were prepared and aliquoted following the manufacturer’s protocol (Goldbio). Luminescence from both luciferases in the same extract was measured separately in different wells, using a 2-second integration time. The translational activity was then calculated as the ratio of Fluc RLU/Rluc RLU as in the mycelium assay. Statistical analyses were performed in R (version 4.4.3) using one-way ANOVA to compare experimental groups. Annotated group means in bar plots indicate statistically homogeneous subsets, such that groups sharing the same letter did not differ significantly at α = 0.05.

## Supplementary Figures and Tables

**Supplementary Table 1.** Accession numbers of all the RNA virus genomes/segments containing self-cleaving ribozymes identified in this study.

EXCEL FILE

**Supplementary Table 2.**
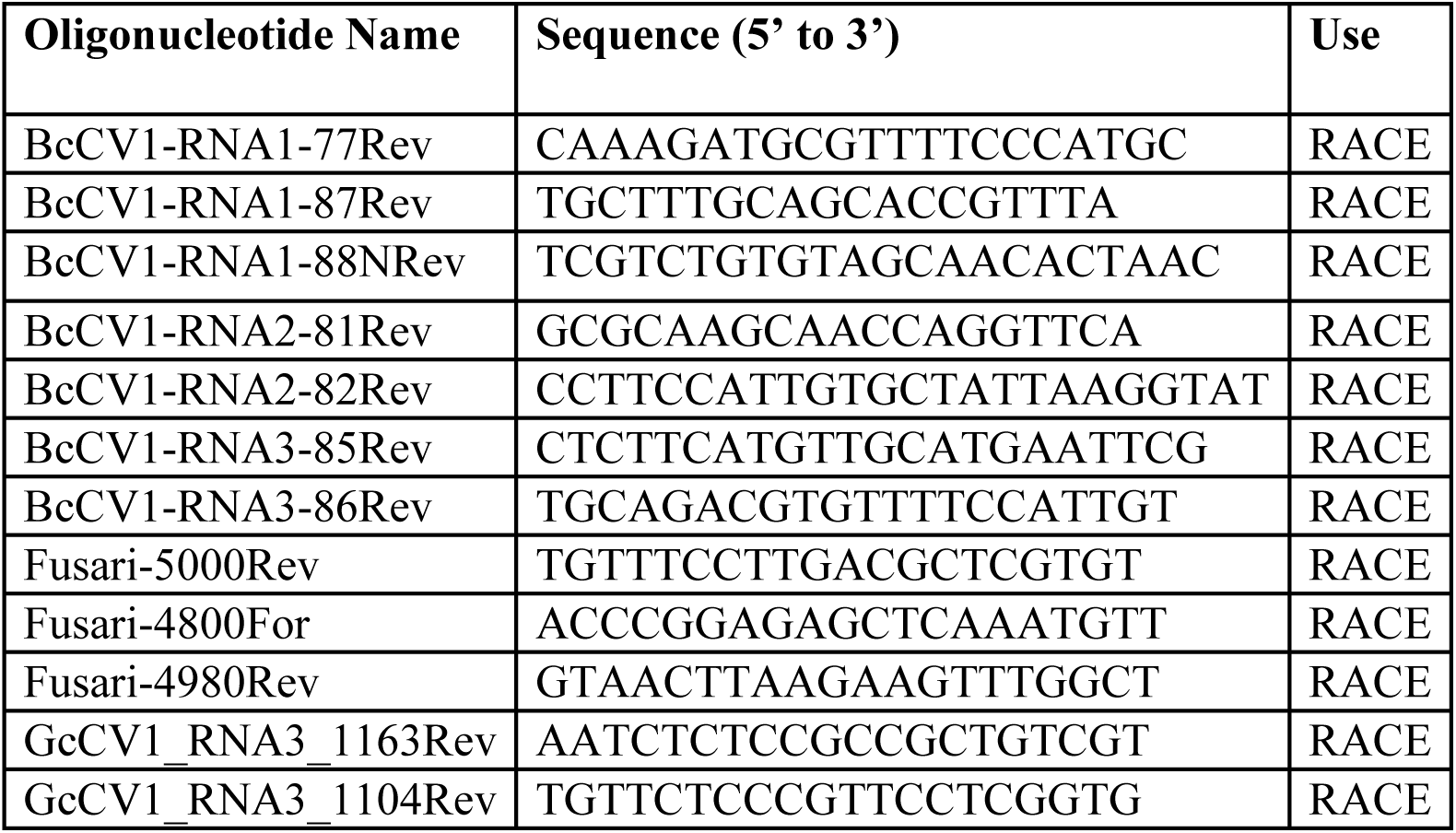
Oligonucleotide sequences and their specific use in this work.

### Supplementary Figures

**Supplementary Fig. 1.**
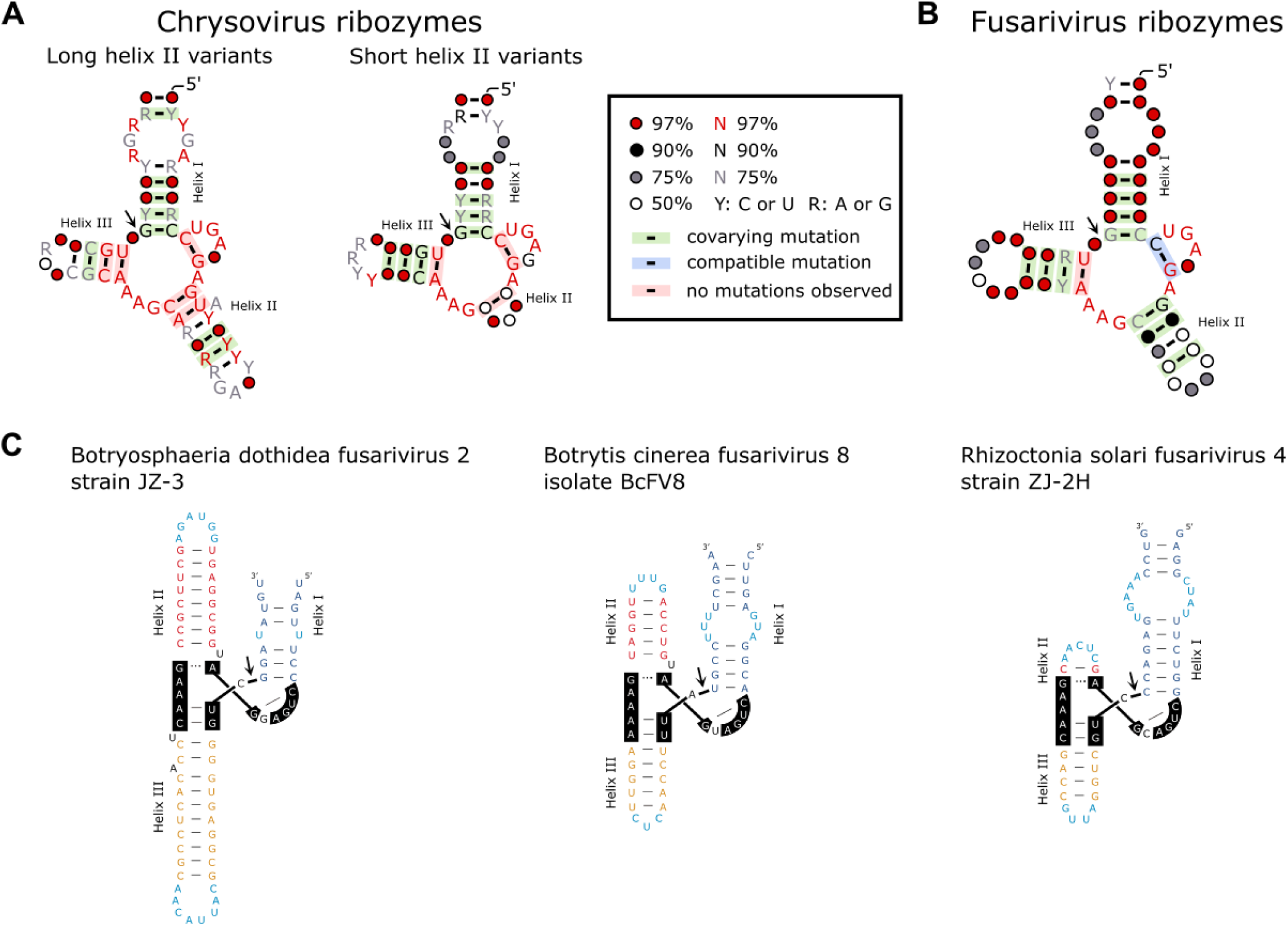
(A) Detailed consensus models and secondary structures of the hammerhead ribozymes either with long helix II (left, based on 26 ribozyme sequences) or short helix II (right, based on 62 ribozyme sequences) from chrysovirus genomes. (B) Consensus model of the hammerhead ribozyme present in most fusarivirus genomes (based on 123 ribozyme sequences). The legend inset applies to the three models shown. (C) Three representative examples of hammerhead ribozymes from fusarivirus genomes with variable stem lengths for helix I, II and III. The site of ribozyme-mediated RNA cleavage is identified by an arrowhead. Consensus models were built as previously described^64^.

**Supplementary Fig. 2.**
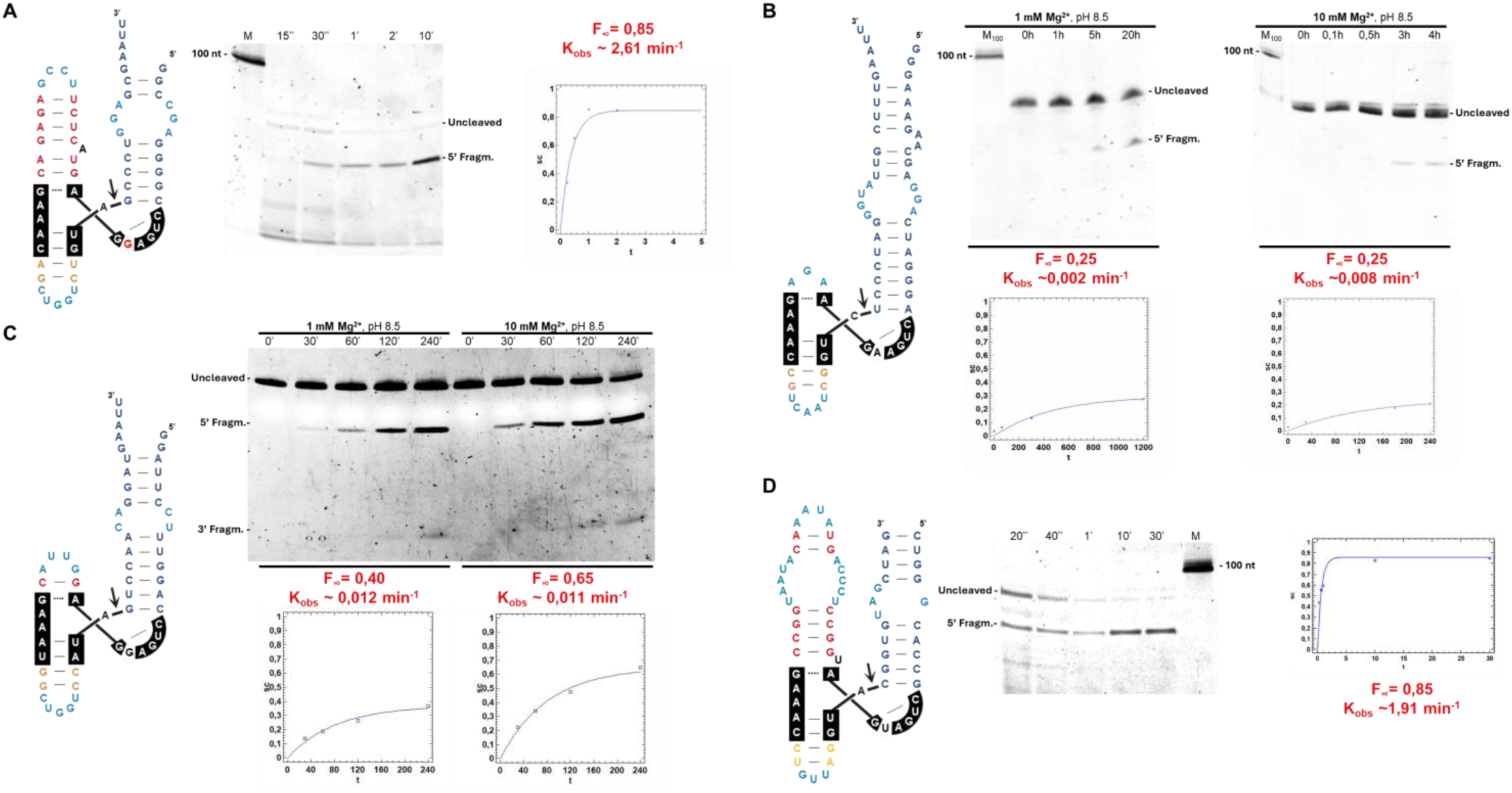
Representative assays of self-cleavage kinetics for four hammerhead ribozymes encoded in fungal and plant RNA viruses. (A) Structure of the hammerhead ribozyme from Verticillium dahliae chrysovirus 1, RNA 1 (left), a 10% PAGE showing the resulting RNAs after different transcription times (standard conditions, pH 7.5, 1 mM Mg^2+^) (center), and the kinetic analysis and quantification graph of the cleavage rates under these co-transcriptional conditions (right). (B) Structure of the hammerhead ribozyme from Brassica campestris chrysovirus (left), two 10% PAGEs showing post-transcriptional assays of self-cleavage with purified RNA of this ribozyme incubated at pH 8.5 and 1 or 10 mM Mg^2+^(Methods) (right), and their kinetic analysis and quantification graph of the cleavage rates (bottom). (C) Structure of the hammerhead ribozyme from Penicillium chrysogenum virus, RNA 1 (left), a 10% PAGE showing a post-transcriptional assay of self-cleavage with purified RNA of this ribozyme incubated at pH 8.5 and 1 or 10 mM Mg^2+^(Methods) (right), and their kinetic analysis and quantification graph of the observed cleavage rates (bottom). (D) Structure of the hammerhead ribozyme from the Rosellinia necatrix megabirnavirus 1 (RnMBV1) (left), a 10% PAGE showing a co-transcriptional self-cleavage assay (center) and quantification graph of the cleavage rate.

**Supplementary Fig. 3.**
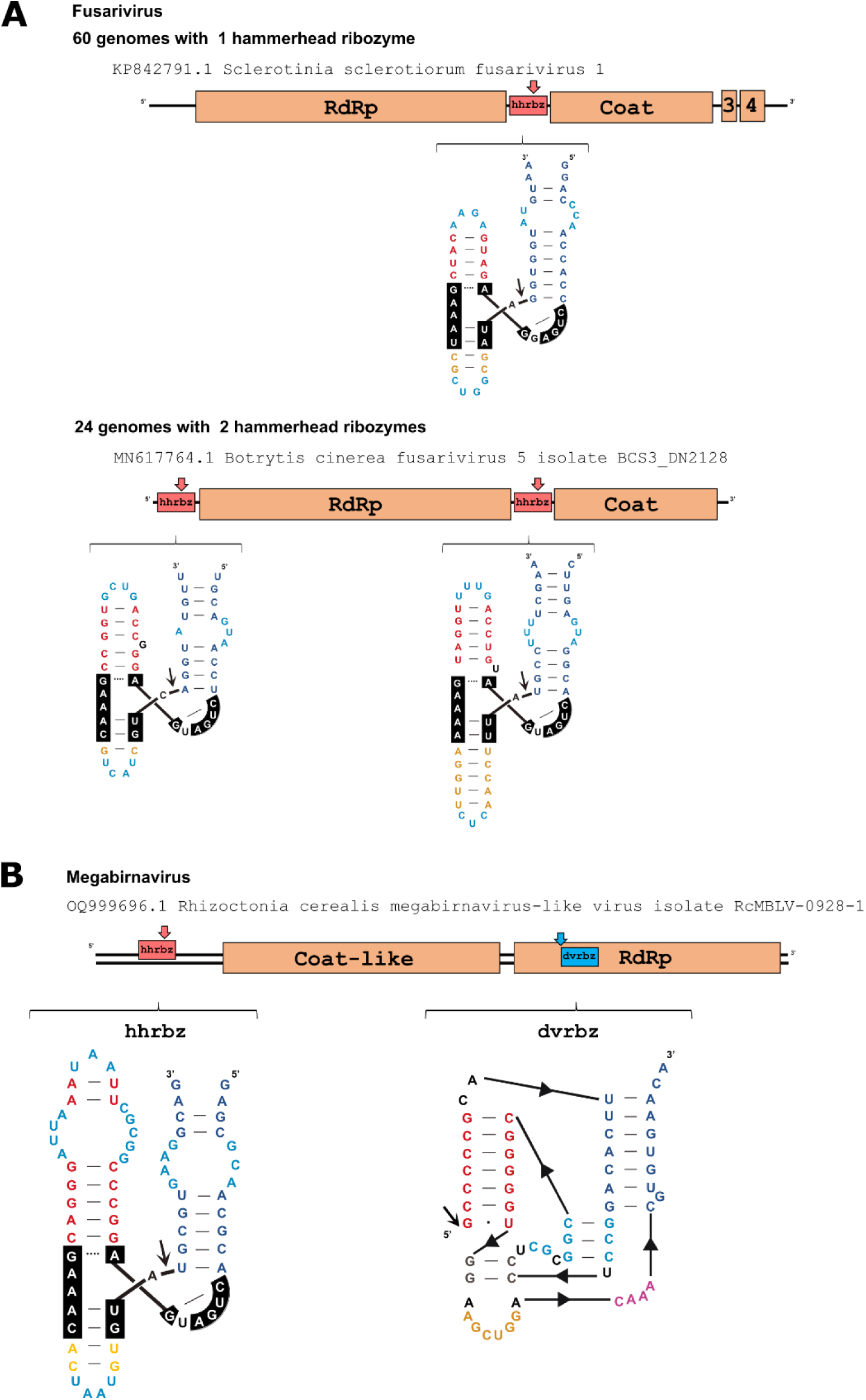
(A) Representative examples of hammerhead ribozymes (hhrbzs) detected in the UTRs of two ssRNA fusariviruses. (B) An example of a dsRNA megabirnavirus-like genome showing the presence of a hhrbz in its 5’-UTR and a deltavirus ribozyme encoded within the predicted RNA dependent RNA polymerase (RdRp) ORF. Sites of ribozyme self-cleavage are indicated by arrows.

**Supplementary Fig. 4.**
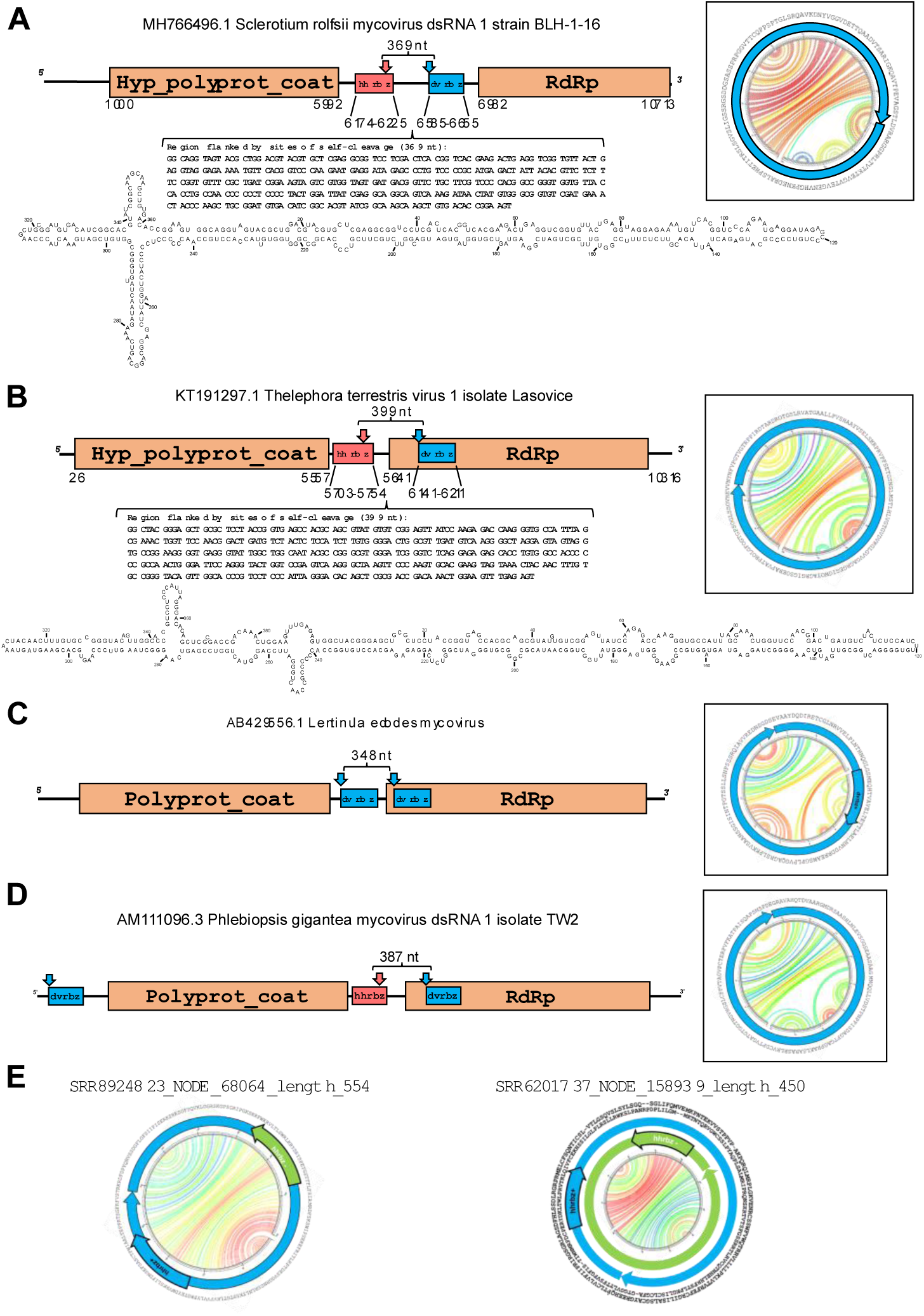
Predicted subgenomic circRNAs detected in (A) Sclerotium rolfsii, (B) Telephora terrestris, (C) Letinula edodes and (D) Phlebiopsis gigantea toti-like mycoviruses containing close ribozyme pairs. Resulting RNAs after double self-cleavage show quasi-rod like structures (circular Jupiter plots are shown at the right), sizes multiple of 3 nt, and absence of stop codons (potentially encoding endless ORFs with protein sequences shown at the insets) as similarly described for zetaviruses^31,32^ (in panel (D), two previously reported Zetavirus genomes are shown for comparison). Hyp; hypothetical. polyprot; polyprotein. hhrbz; hammerhead ribozyme. dvrbz; deltavirus ribozyme. RdRp; RNA dependent RNA polymerase.

**Supplementary Fig. 5.**
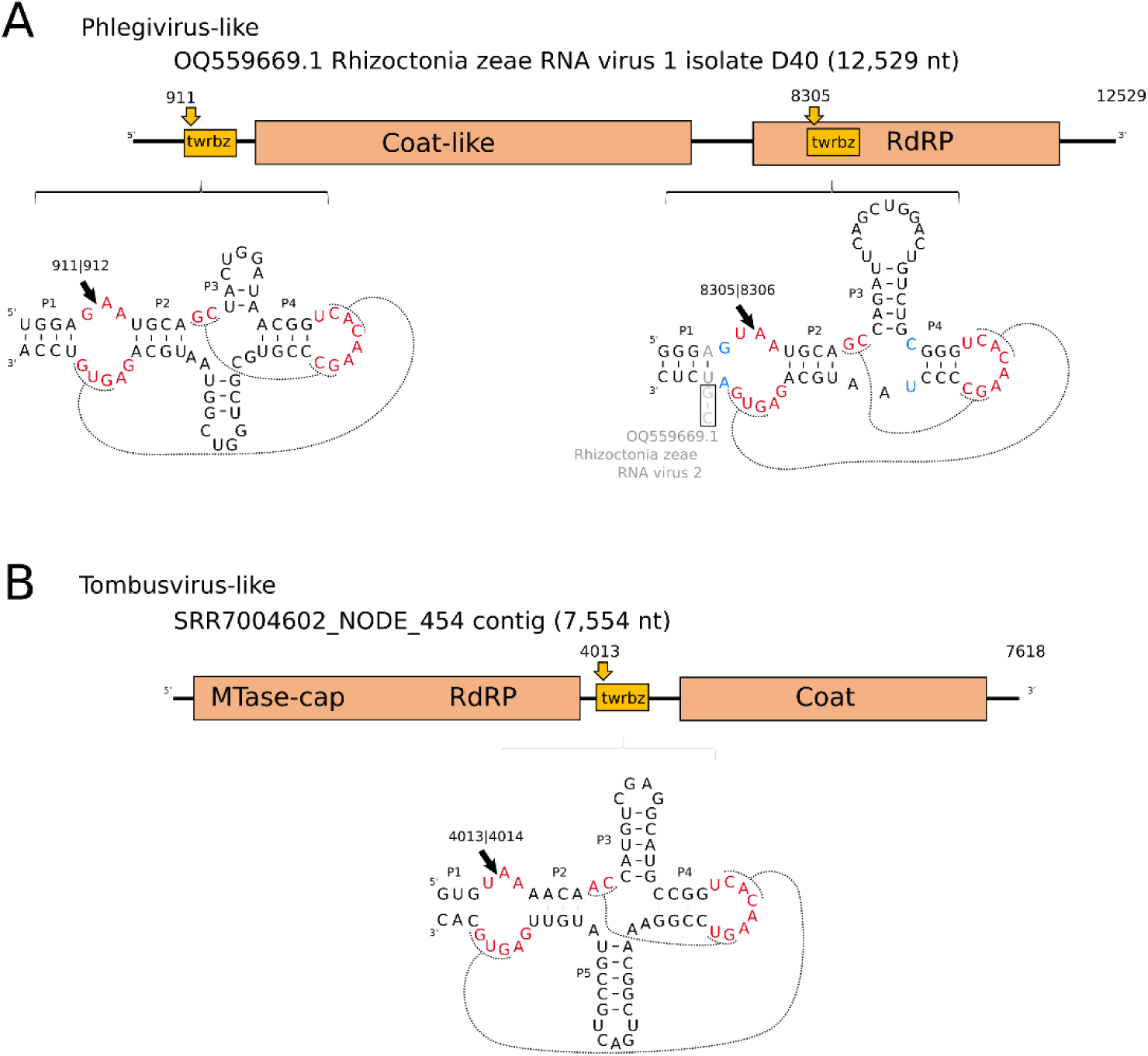
Twister ribozymes (twrbz) motifs detected in the RNA genomes of (A) phelgivirus-like and (B) tombusvirus-like sequences. Highly conserved nucleotides and unexpected changes in the consensus structure of most twrbz motifs are depicted in red and blue, respectively.

**Supplementary Fig. 6.**
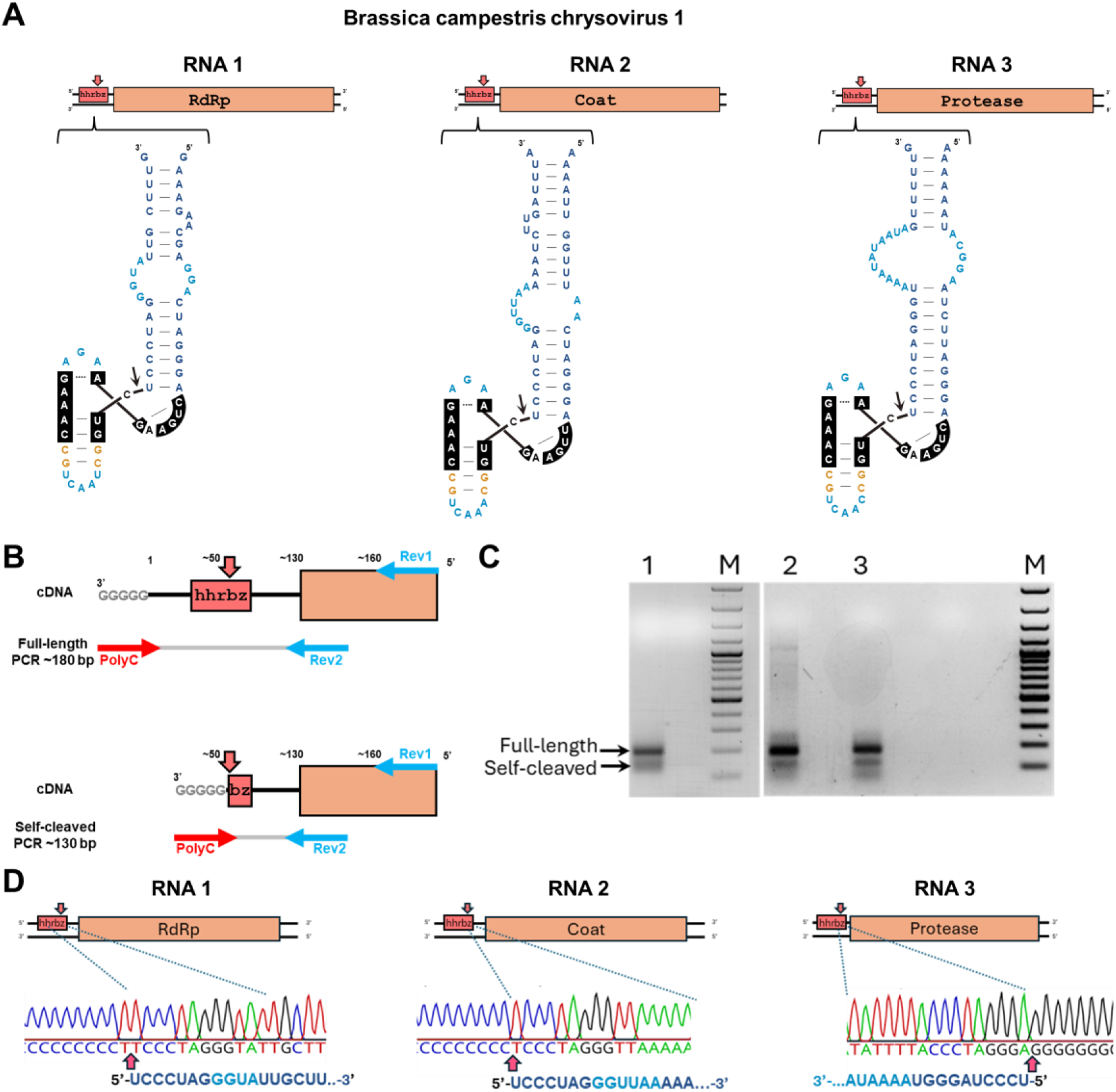
5’ RACE of Brassica campestris chrysovirus 1 (BcCV-1) from infected cabbage Torzella plants. (A) Schematic representation of the BcCV-1 hhrbzs in the 5’-UTRs of each of the three genomic RNA segments. (B) Schematic representation of the aproximated locations of the oligos used for the RACE experiments and the expected sizes of the PCR products resulting from the full-length and self-cleaved RNAs (C) Agarose gel analysis of 5’ RACE PCR of BcCV-1 RNA1, RNA2 and RNA3 (lanes 1, 2 and 3, respectively) shows two bands corresponding to the amplification of the full-length (upper and more abundant band) and the self-cleaved (lower band) RNAs. For the RNA1 (lane 1), a nested PCR, using a reverse BcCV-1 RNA1 (BcCV1-RNA1-88NRev, Supplementary Table S2), nested to the primer used in the first RACE amplification (BcCV1-RNA1-87Rev, Supplementary Table S2), is shown. M, DNA molecular weight marker (100 bp DNA ladder, Thermo Fisher). (C) Sequencing electropherograms of the cloned RACE product of smaller size for each RNA genomic segment. The 5’ end coincides with that predicted to be generated by the hhrbz self-cleavage (indicated by arrows).

**Supplementary Fig. 7.**
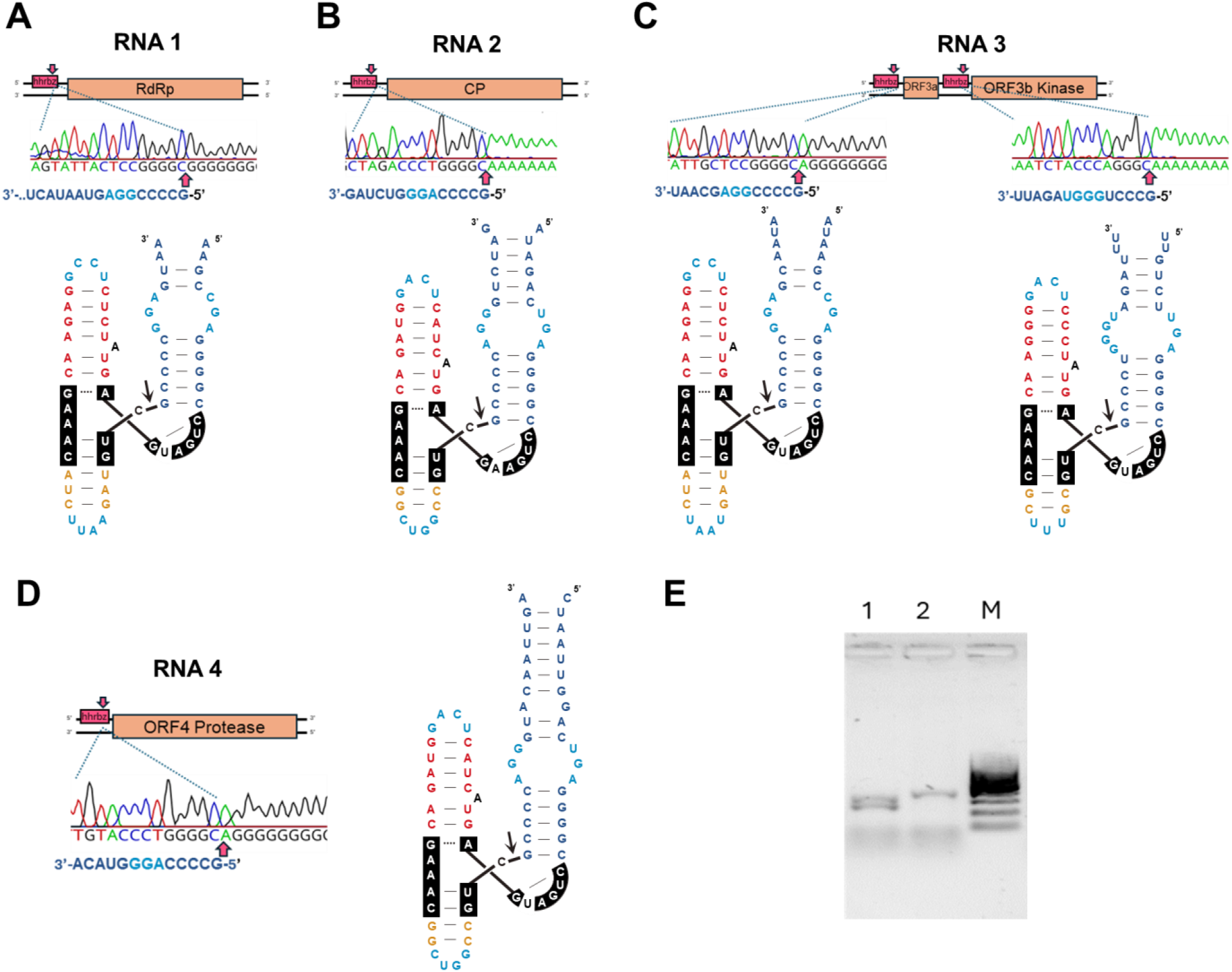
5’ RACE of Gnomoniopsis castaneae chrysovirus 1 (GcCV-1) from infected *G. castaneae* isolate 1^44^. Schematic representation of GcCV-1 RNAs, hammerhead ribozymes (hhrbz) secondary structure and sequencing electropherograms of the cloned RACE product of smaller size for each (A) RNA 1, (B) RNA 2 (C) RNA 3 and (D) RNA 4 genomic segments. Hhrbzs are present in all the 5’-UTR of RNAs, whereas RNA3 shows a second intergenic hhrbz motif between the two conserved ORFs (3a and 3b). The 5’ ends of the sequencing electropherograms coincide with those predicted to be generated by the hhrbz self-cleavage (indicated by arrows). (E) Agarose gel analysis of 5’ RACE PCR (lane 1) and 3’RACE PCR (lane 2) of GcCV-1 RNA2. Lane 1 shows two bands corresponding to the amplification of the full-length (upper band) and the self-cleaved (lower band) RNAs. M; DNA molecular weight marker of 100 bp DNA (Thermo Fisher). RdRp; RNA dependent RNA polymerase. CP; coat protein. ORF; open reading frame.

**Supplementary Fig. 8.**
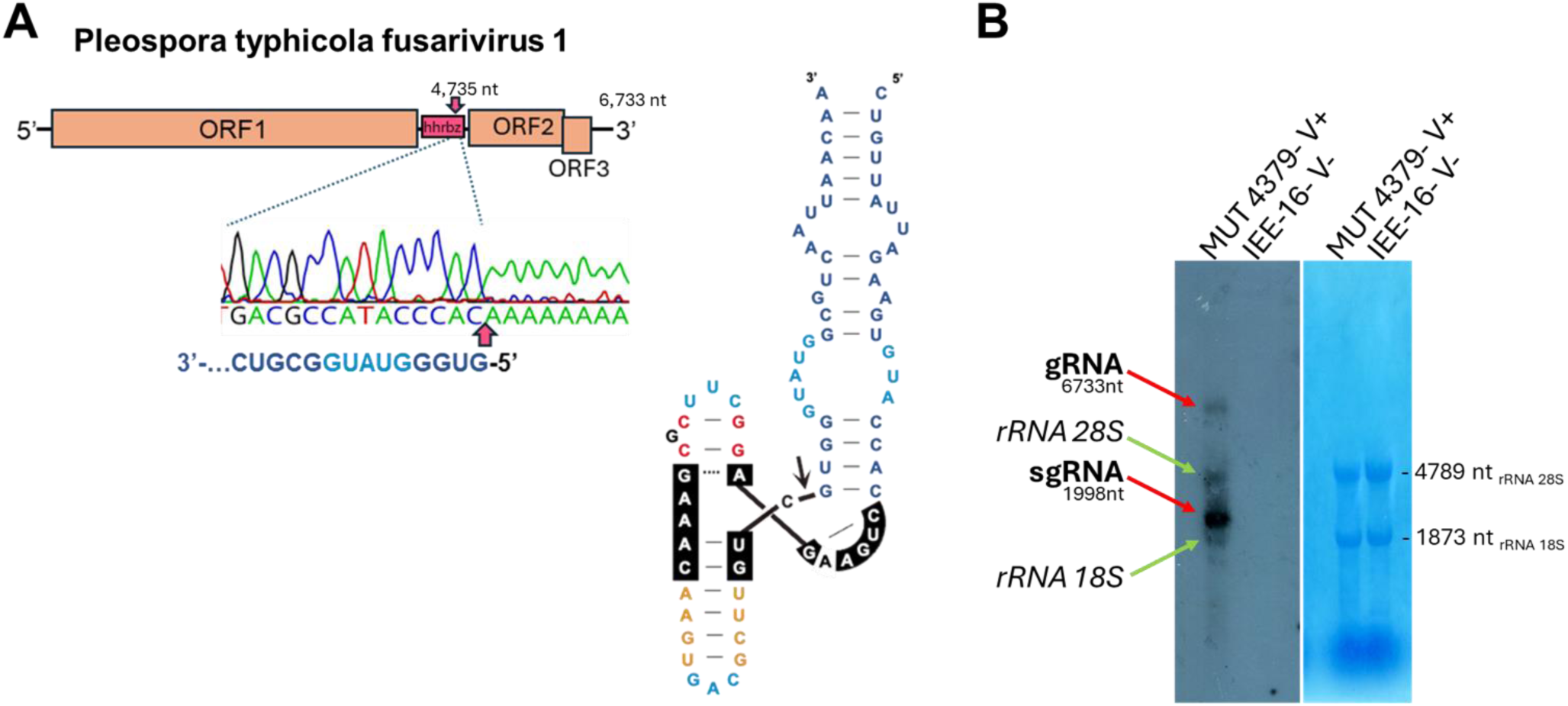
5’ RACE of Pleospora typhicola fusarivirus 1 (PtFV1) from infected MUT 4379 ^45^. (A) Genome organization, with the position of the hammerhead ribozyme (hhrbz) placed in the intergenic region. The secondary structure of the ribozyme is also displayed (right), as well as the sequence electropherogram showing the 5’ end of the cleavage site (red arrow). (B) Northern blot analysis of RNA extracts from virus-infected isolate MUT4379 and virus-free *Rhizoctonia solani* isolate IEE-16^73^. On the left panel, red arrows indicate the PtFV1 genomic RNA (gRNA) and PtFV1 sgRNA positions, while green arrows indicate some possible cross-hybridization with the two fungal ribosomal RNAs. The right panel shows a methylene blue stain of the membrane displaying the two ribosomal RNA bands.

**Supplementary Fig. 9.**
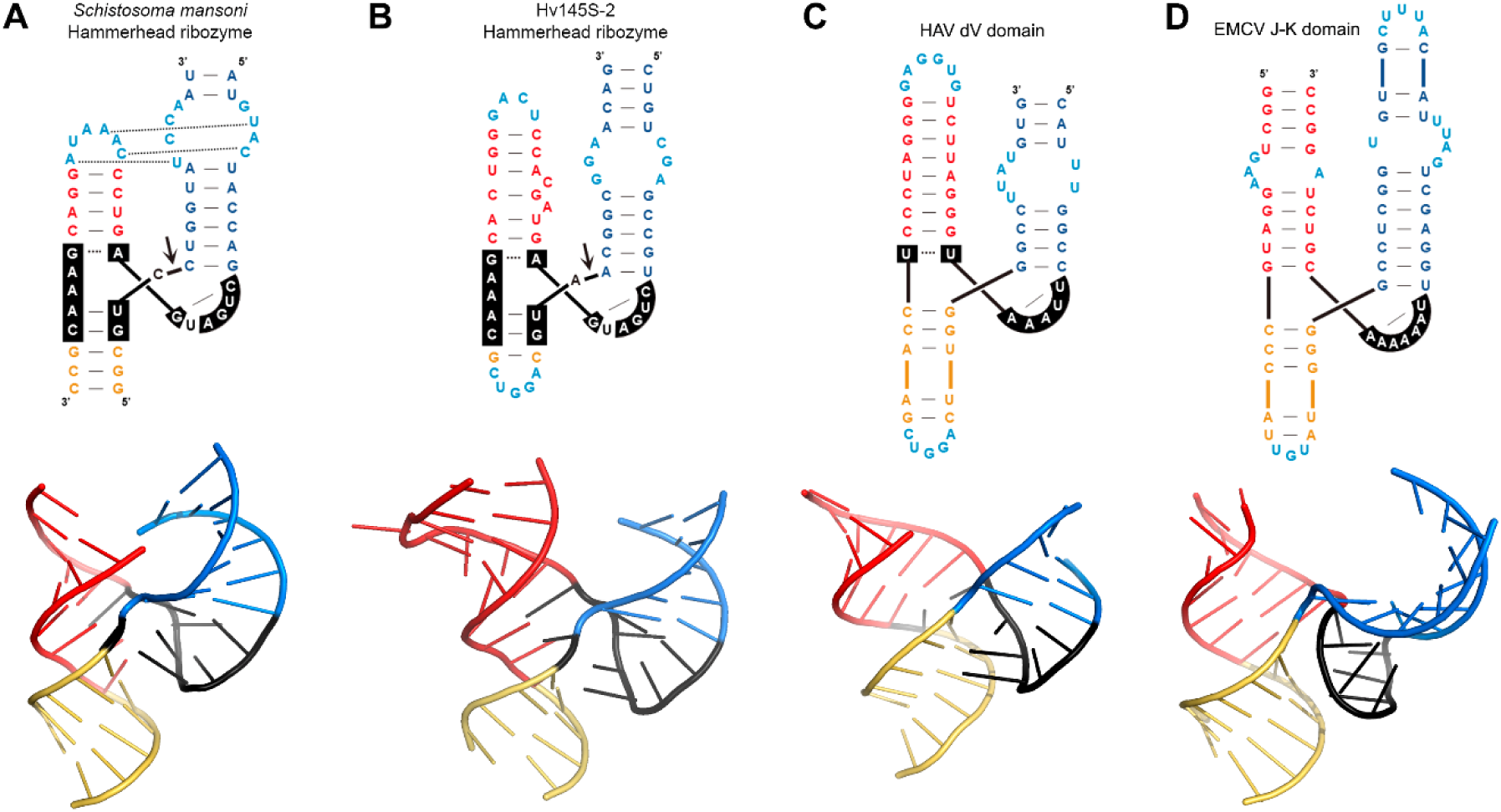
(A) Top: schematic representation of a type I hammerhead ribozyme (hhrbz) from Schistosoma mansoni with tertiary loop-loop interactions indicated by dotted lines. Bottom: 3D-structure (pdb 3ZD5) showing the central conserved core in black and the three stems I, II, and III in blue, red, and yellow, respectively; loops are not shown. Similarly, schematic (top) and 3D structures (bottom) of (B) the Hv145S-RNA2 hhrbz (predicted through the Alphafold 3 server^74^), (C) Hepatitis A virus (HAV) IRES domain V (pdb 6mwn)^61^ and (D) the J-K domain of the encephalomyocarditis virus (EMCV) IRES element (pdb 2nbx)^59^.

**Supplementary Fig. 10.**
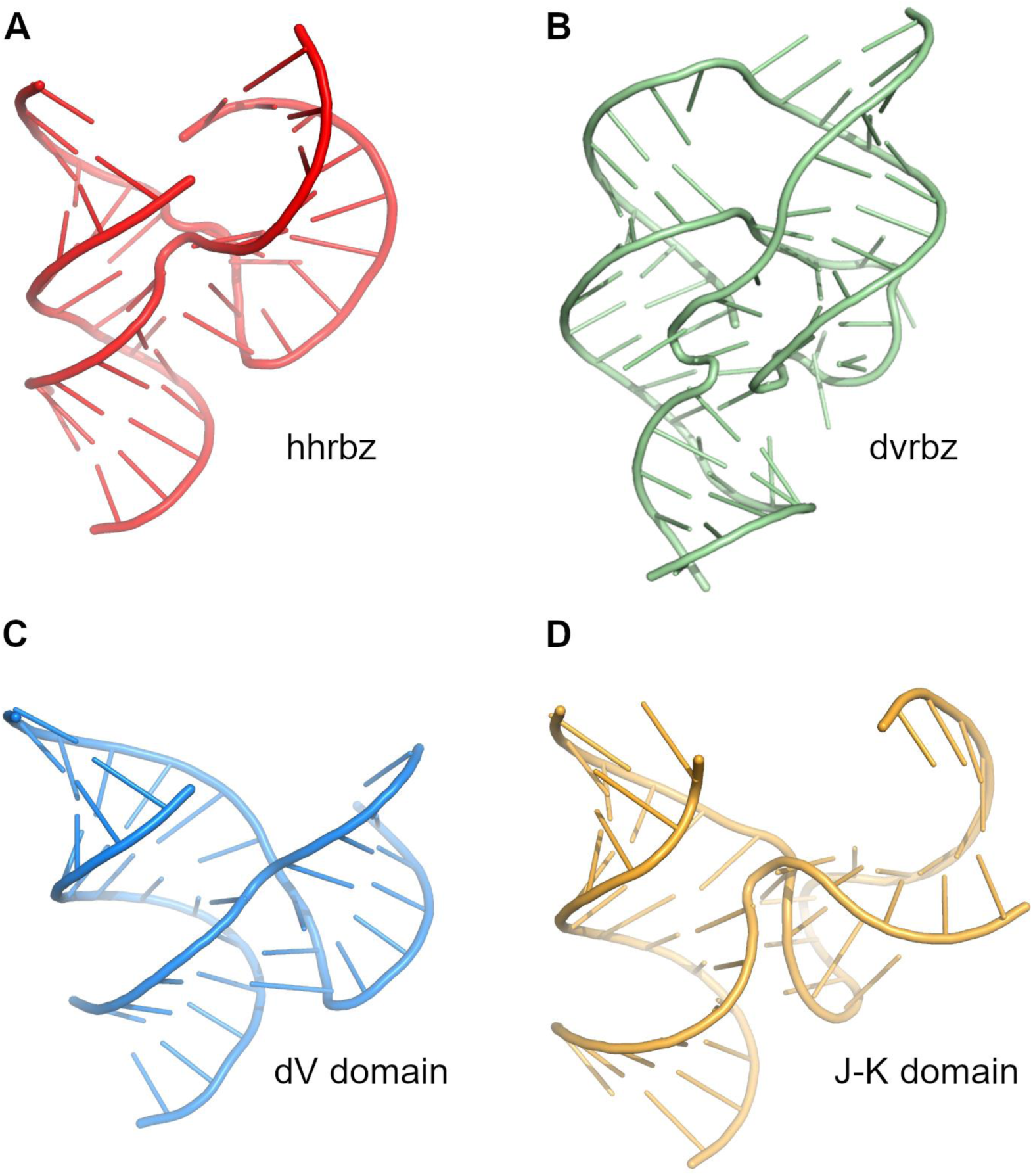
Three dimensional models for (A) the hammerhead ribozyme from *Schistosoma mansoni* retrozymes (pdb 3ZD5, central core and stems, but not interacting loops 1 and 2, are shown), (B) the human Hepatitis Delta ribozyme (pdb 1DRZ), (C) the Hepatitis A virus IRES dV core domain (pdb 6mwn) and (D) the encephalomyocarditis virus IRES J-K core domain (pdb 2nbx).

## Acknowledgments

This work was funded by: University of Valencia Margarita Salas Fellowship MS21-067 (MJLG); Generalitat Valenciana Grant PROMETEO CIPROM/2022/21 (MdlP); Ministerio de Ciencia, Innovacion y Universidades de España-FEDER grant PID2024-162245NB-I00 (MdlP). We are grateful to Daniela Alio and Maria Minutolo for providing plants infected by Brassica campestris chrysovirus 1.

